# Mechanistic target of rapamycin (mTOR) regulates self-sustained quiescence, tumor indolence and late clinical metastasis in a Beclin-1-dependent manner

**DOI:** 10.1101/2022.05.05.490726

**Authors:** Carole Nicco, Marine Thomas, Julie Guillermet, Maryline Havard, Fanny Laurent-Tchenio, Ludivine Doridot, François Dautry, Frédéric Batteux, Thierry Tchenio

## Abstract

Self-sustained quiescence (SSQ) has been characterized as a stable but reversible non-proliferative cellular state that limits the cloning of cultured cancer cells. By developing refined clonogenic assays, we showed here that cancer cells in SSQ can be selected with anticancer agents and that culture at low cell density induced SSQ in pancreas and prostate adenocarcinoma cells. Pre-culture of cells in 3D or their pretreatment with pharmacological inhibitors of mechanistic target of rapamycin (mTOR) synergize with low cell density for induction of SSQ in a Beclin-1-dependent manner. Dissociated pancreatic adenocarcinoma (PAAD) cells rendered defective for SSQ by down-regulating Beclin-1 expression exhibit higher tumor growth rate when injected subcutaneously into mice. Conversely, dissociated PAAD cells in SSQ promote the formation of small indolent tumors that eventually transitioned to a rapid growth phase*. Ex vivo* clonogenic assays showed that up to 40% of clonogenic cancer cells enzymatically dissociated from resected fast-growing tumors could enter SSQ, suggesting that SSQ could significantly impact the proliferation of cancer cells that are naturally dispersed from tumors. Remarkably, the kinetics of clinical metastatic recurrence in 124 patients with pancreatic adenocarcinoma included in the TGCA-PAAD project could be predicted from Beclin-1 and Cyclin-A2 mRNA levels in their primary tumor, Cyclin A2 mRNA being a marker of both cell proliferation and mTOR complex 1 activity. Overall, our data show that SSQ is likely to promote the late development of clinical metastases and suggest that identifying new agents targeting cancer cells in SSQ could help improve patient survival.

## Introduction

Metastases are the leading cause of cancer-related morbidity and mortality and account for more than 80% of cancer deaths. Metastasis involves cancer cells, isolated or in clusters of a few cells, that are disseminated from the primary tumors to distant sites by lymphatic or hematogenous routes. Metastases development is an inefficient process. In experimental models, only a small fraction of cancer cells that have extravasated in the parenchyma of a distant organ begins to proliferate to form micro-metastases, with most of cancer cells entering into a non-growing state called dormant ^1–3^. Additionally, only a small subset of micro-metastases continue to grow to form macroscopic tumors ^1–3^ and the growth of small macroscopic tumors into larger tumors is also limited ^4, 5^. Unfortunately, the mechanisms that limit development of metastases are also probably involved in the persistence of cancer cells in patients in complete remission and are a cause of the late development of metastases ^5–7^.

Since a metastase is basically a clonal cell population resulting from the proliferation of a single disseminated cancer cell, we have undertaken studies on the factors limiting the cloning of cancer cells in culture in order to better understand the factors regulating the development of metastases. This led us to discover that the ability of prostate adenocarcinoma (PA) cells to proliferate at low cell density to form a cell clone can be severely limited by a phenomenon we initially called dormancy ^8, 9^, by analogy with the phenomenon of cancer cell dormancy ^5^, then renamed Self-Sustained Quiescence (SSQ) ^10^ to highlight its cell-autonomous feature. Indeed, PA cells can be induced to enter a G0/G1 growth-arrested state that, once established, maintains itself at low cell density despite the return of cells to growth-permissive conditions. We showed that this self-sustained growth arrest involves a stable decrease of the redox potential of cellular thiols and is further stabilized by BMP signaling pathways ^10^. Accordingly, cells in SSQ can be switched to a proliferating state when treated with SSQ inhibitors (SSQi) consisting of a reduced thiol (such as dithiothreitol, glutathione or N-acetylcysteine) and/or an inhibitor of BMP signaling (such as K02288; see also the Methods section for more details) ^8–10^. Additionally, SSQ can be destabilized by autocrine factors of less than 2 kDa produced by most types of cancer cells grown in 2D culture (see Fig S1 in ^9^). The self-maintenance property of SSQ is remarkable compared to other models of cancer cell dormancy that in most cases involved mechanisms that are non-cell-autonomous in accordance with the “seed and soil” model. This raises the issue of the biological relevance of SSQ in the regulation of tumor development. Here we showed that PA and pancreatic ductal adenocarcinoma (PADC) cells in SSQ or competent for SSQ can be selected with anticancer agents after subculturing cells at low cell density. This allowed the characterization of factors modulating induction of SSQ and the isolation of PDAC cell populations highly enriched in SSQ that we used for *in vivo* studies. We then applied the results obtained to the analysis of metastasis in pancreas adenocarcinoma (PAAD) patients.

## Results

### Selection of clonogenic cancer cells in SSQ with anticancer agents at low cell density

We hypothesized that anticancer agents that preferentially target proliferative cells could select cells in SSQ. We used KC-DT66066 cells, derived from a pancreas ductal adenocarcinoma (PDAC) developed in a genetically engineered LSL-Kras^G12D/+^, Pdx1-Cre mouse model of spontaneous PDAC ^11, 12^ and routinely passaged at medium/high cell density in 2D-culture plates. When these cells were plated at low cell density in regular growth medium, they exhibited high cloning efficiency (CE=48 ± 14 % of seeded cells grew to form cell clones; mean ± sd, n=24; see the none⟶ none clonogenic assay in Fig 1A-B) that was slightly increased by addition of SSQi (59 ± 17 %; see the none⟶ SSQi clonogenic assay in Fig 1A-B). To test for the selection of cancer cells in SSQ with gemcitabine, KC-DT66066 cells were plated at low density in regular growth medium and cultured for 3 days before applying gemcitabine selection for 4 days. When cells were washed and returned to their regular growth medium, very few, if any, growing cell clones were detected after 10-12 days, even after microscopic examination (see gem d3⟶ none in Fig 1A-B). However, when cells were returned to regular growth medium supplemented with SSQi after gemcitabine selection, growing cell clones were recovered at an averaged frequency of 1.5% of seeded cells (see gem d3⟶ SSQi in Fig 1A-B). This showed that non-growing cells that had survived to gemcitabine treatment were in SSQ since they switched to a proliferative state only after adding SSQi. Interestingly, the recovery of growing cell clones was reduced to background (<0.4%) if gemcitabine was added immediately after cell seeding (see gem d0⟶ SSQi in Fig 1A-B). This indicated that most cells from 2D-cultures entered SSQ only after plating at low cell density.

**Fig 1:**
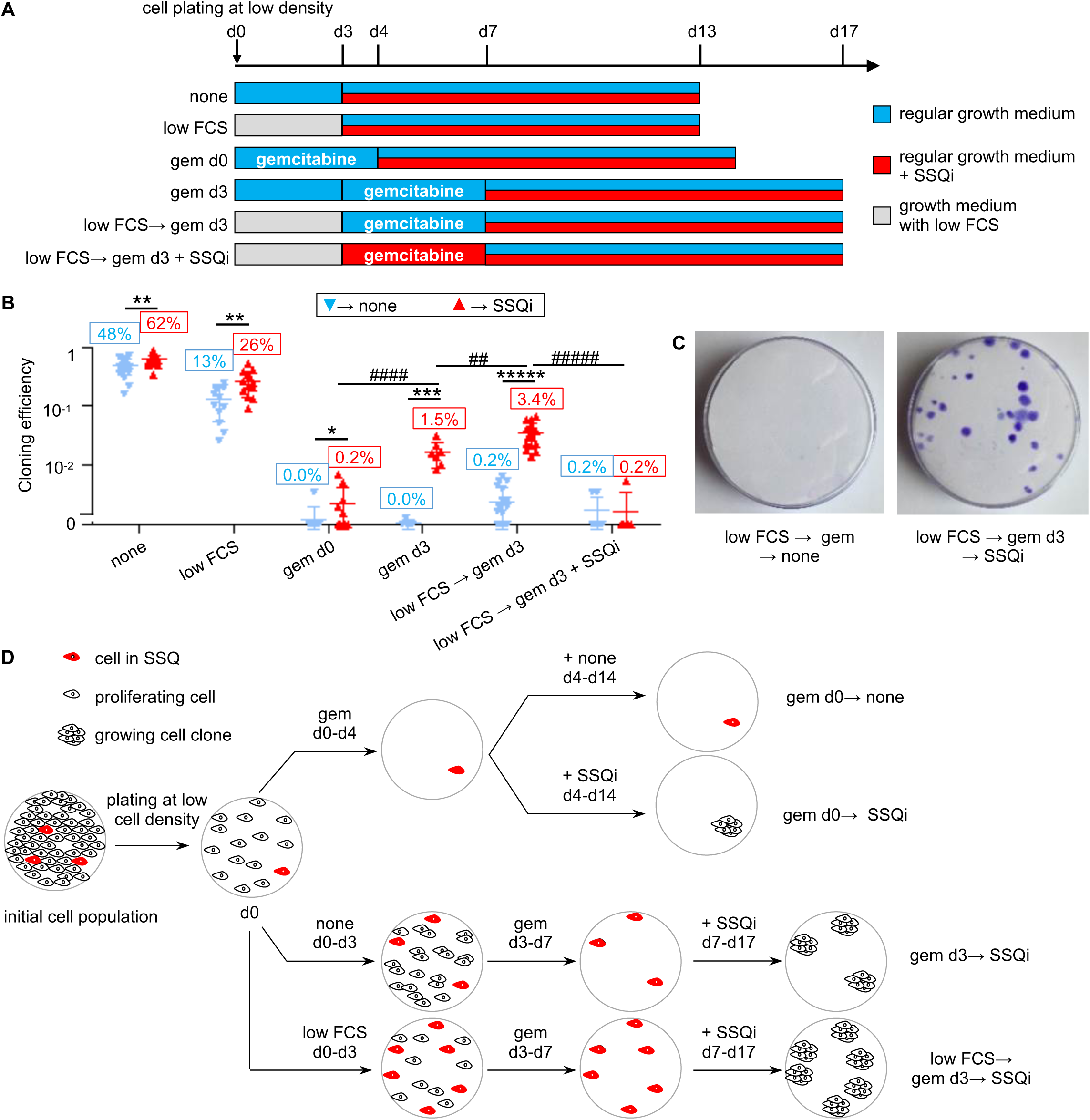
Selection of pancreas cancer KC-DT66066 cells in SSQ at low cell density with anticancer agents. **A)** Outlines of clonogenic assays to assess the kinetics of SSQ formation. Cells were plated at low cell density on day 0 (d0) and cultured as indicated. The labels on the left are those used in the scatter plots. SSQi is a mixture of pharmacological agents that reverts SSQ (see Methods). **B)** Kinetics of the generation of gemcitabine-resistant cells. Scatter plots of cloning efficiency (CE) measured with the indicated clonogenic assays. Labels in the red boxes indicate the percentage of seeded cells having formed a cell clone. * indicates the statistical significance of the change in CE induced by SSQi supplementation and # the statistical significance of the difference in CE between the indicated clonogenic assays with SSQi supplementation. **C)** Representative pictures of fixed and stained cell culture dishes after culturing the cells as indicated. **D)** Rationale for determining the initial frequency of cancer cells in SSQ in a given cell population and the frequency at which they are induced by cell subculture at low cell density or low cell density and low serum. The gem d0 clonogenic assay selects cells that were already in SSQ in the initial cell population. The selected SSQ cells switch to a proliferating state and formed cell clones after the addition of SSQi (gem d0⟶ SSQi clonogenic assay). Growing cell clones obtained in clonogenic assays with delayed gemcitabine selection and SSQi supplementation (gem d3⟶ SSQi and low FCS⟶ gem d3⟶ SSQi) originated from cells already in SSQ in the initial population or that were induced to enter SSQ by subculture at low cell density or at low cell density and in low serum.

As this induction of SSQ could be counteracted by growth factors that promote cell proliferation, we tested whether low fetal calf serum concentration in growth medium (low FCS) could facilitate SSQ induction after plating at low cell density. After plating for 3 days at low cell density in low FCS, spontaneous cloning efficiency (CE) decreased to 13% of seeded cells but was increased twofold by adding SSQi (see low FCS in Fig 1A-B). This suggests that clonogenic cancer cells that survive to culture at low cell density in low serum were more susceptible to the stimulation of clonal cell growth by SSQi. Correspondingly, frequency of cancer cells in SSQ selected by gemcitabine increased to 3.4% of seeded cells under these conditions, showing that reduced serum concentration synergized with low cell density to induce SSQ (see low FCS⟶ gem d3⟶ SSQi in Fig 1A-C).

Of note, the addition of SSQi could be delayed for at least 3 days after completion of gemcitabine selection with no significant decrease of cell clone recovery, showing that SSQi did not rescue cancer cells undergoing a delayed cell death phenomenon that might have been induced by gemcitabine treatment (Fig S1A). In fact, the addition of SSQi at the same time as gemcitabine completely blunted the subsequent recovery of growing cell clones (see low FCS⟶ gem d3 + SSQi⟶ SSQi in Fig 1A-B). This showed that SSQi sensitized cells to gemcitabine, as expected if the main action of SSQi is to promote the switch from SSQ to a proliferative state. These observations were generalized to others pancreas and prostate cancer cell lines (Fig S1B, D-E) and anticancer drugs exhibiting a strong and selective toxicity to proliferating cancer cells (Fig S1D-E). Finally, we showed that SSQ entry and exit are fully reversible processes that can be repeated serially since proliferating cell populations derived from cancer cells in SSQ selected with nocodazole or gemcitabine and reactivated with SSQi could re-enter in SSQ with high efficiency when cultured at low cell density (Fig S1F). In the remainder of this article, the clonogenic assay gem d0⟶ SSQi was used to measure the relative frequencies of cells already in SSQ in the cell populations studied (Fig 1D). The relative frequency of clonogenic cancer cell in SSQ measured by clonogenic assays with delayed selection by gemcitabine (gem d3⟶ SSQi or low FCS⟶ gem d3⟶ SSQi) corresponds to the sum of the relative frequency of cancer cells already in SSQ in the cell population studied and the relative frequency of their induction by low cell density and low serum (Fig 1D).

### SSQ is promoted by 3D cell culture and treatment with pharmacological inhibitors of mTOR

We observed that KC-A338 cells entered SSQ at high frequency in 2D culture (Fig S1B), which correlated with their ability to spontaneously form transformed foci that eventually detached from the bottom of the culture dish to form compact tumor spheroid-like structures (Fig S1C). This led us to investigate whether 3D culture could promote SSQ. 3D culture of KC-DT66066 cells on cell-surface-repellent plates in their regular growth medium resulted in the formation of small, non-growing and loosely attached cell aggregates (Fig 2A-B). However, when growth medium was supplemented with conditioned medium (CM) harvested from senescent mouse fibroblasts cultures that produce numerous growth factors and cytokines ^13^, compact growing spheroids were obtained (Fig 2A-B). Analysis of the expression of stemness markers by RT-qPCR showed that non-growing cell aggregates overexpressed *Aldh1* while growing spheroids overexpressed *Lgr5* (Fig 2C and see the S1 file for the full list of genes tested). Interestingly, the gem d0 ⟶ SSQi clonogenic assay showed that 6-7 % of clonogenic cancer cells had entered SSQ during 3D culture both as non-growing cell aggregates or growing spheroids (Fig 2D). Moreover, cells from 3D culture exhibited an increased rate of SSQ induction after plating at low density in reduced serum (compare low FCS⟶ gem d3 ⟶ SSQi between 2D control, 3D and 3D+CM in Fig 2D). In an attempt to understand why SSQ was promoted by 3D cell culture, we examined the possible role of autophagy since cell detachment from the extracellular matrix can induce autophagy ^14^. Therefore, we used the autophagy gene reporter GFP-LC3-RFP-LC31′G ^15^ to measure the autophagic flux and observed that it was stably enhanced in 3D cultured cells (Fig 2E-G). This led us to examine whether stimulation of autophagy in 2D cell culture could promote SSQ. Accordingly, cells grown at medium cell density in standard culture plates were pretreated for 1 day with Torin1 ^16^, a standard inhibitor of mTOR that strongly induces autophagy ^15^, then harvested and seeded at low cell density to test for SSQ induction as described in Fig 1A. Torin1 pretreatment strongly increased the induction of SSQ by plating cells at low density in reduced serum (compare low FCS⟶ gem d3 ⟶ SSQi between pretreated and non-pretreated cells in Fig 2D). Moreover, the gem d0⟶ SSQi clonogenic assay indicated that at least 2 % of clonogenic cancer cells were induced to enter SSQ before plating cells at low density. These data were confirmed using rapamycin, the canonical mTOR inhibitor and both Torin1 and rapamycin also increased SSQ in human prostate adenocarcinoma cell lines (Fig S2A-B). Taken together, these results showed that inhibition of mTOR signaling pathways was sufficient to strongly promote the induction of SSQ in cancer cells.

**Fig 2:**
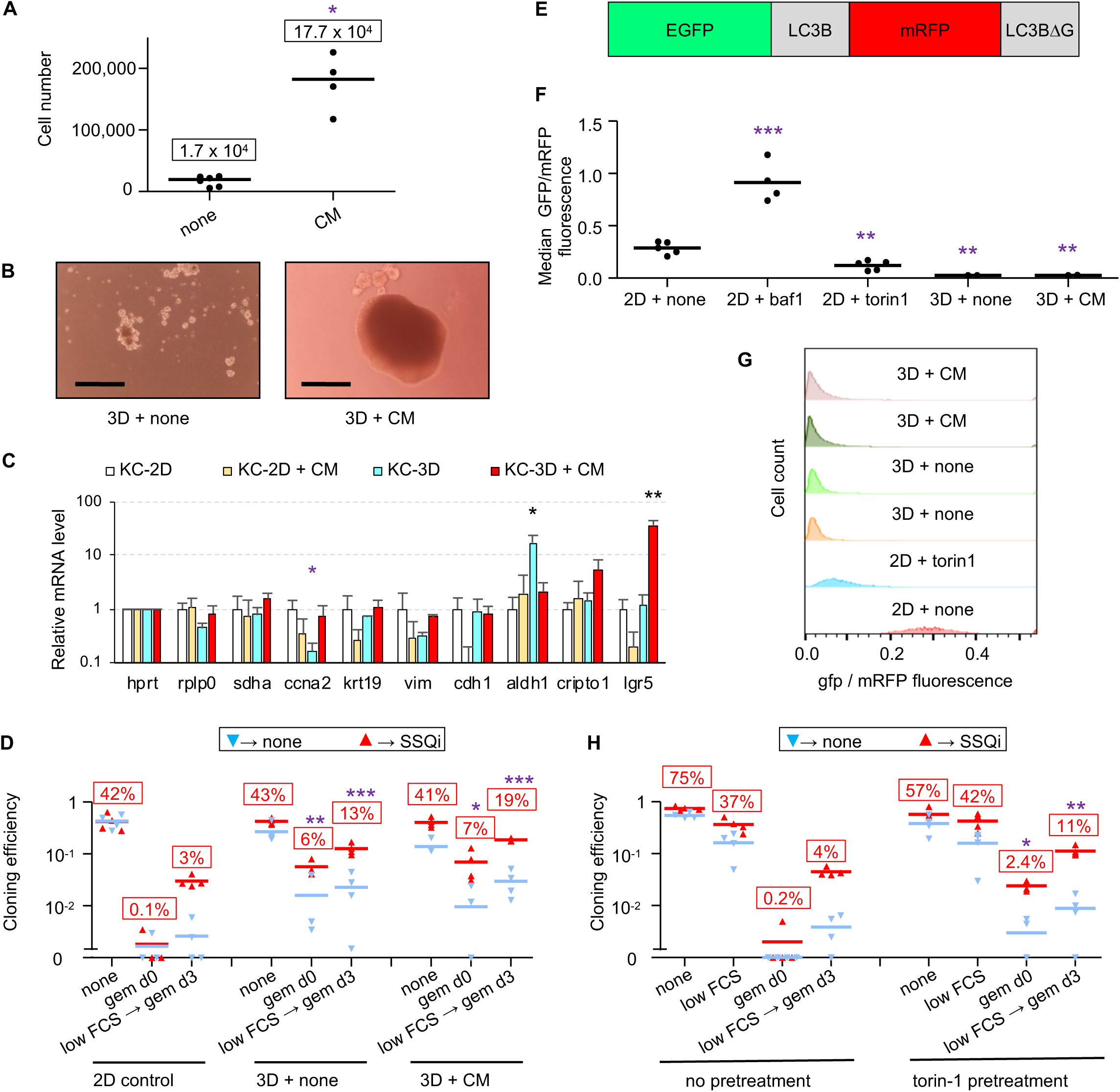
SSQ induction in KC-DT66066 cells is enhanced by 3D cell preculture or pretreatment with pharmacological inhibitors of mTOR in 2D cell cultures. **A)** Growth of cells seeded at 10^4^ cells per well in cell-repellent surface 12-well plate and cultured for 10 days in regular growth medium supplemented with none (none) and 50% conditioned medium (CM) from senescent mouse embryo fibroblasts cultures. The stars indicate the statistical significance of the differences with cells grown in regular growth medium. Labels indicate mean cell number. **B)** Representative pictures of cells grown for 7 days in cell-repellent surface plates in regular growth medium supplemented with none (3D + none) or CM (3D + CM). The scale bar indicates 0.25 mm. **C)** RT-qPCR analysis of mRNA species levels according to the indicated cell-culture conditions. Stars indicate statistical significance of differences with the KC-2D condition. 4 independent experiments were performed. **D)** Clonogenic assays to compare SSQ induction in cells from 3D or 2D cultures using the same cell dissociation method. Note that the relatively high frequency of cell clones in the clonogenic assays gem d0→ none and low FCS→ gem d3→ none resulted from the spontaneous reversion of cells in SSQ after gemcitabine selection when their frequency exceeded 10% of the seeded cells. **E)** Schematic representation of the fusion protein encoded by the reporter gene EGFP-LC3B-mRFP-LC3B1′G encoding the autophagy sensitive EGFP-LC3B and the autophagy resistant mRFP-LC31′G fluorescent proteins. **F)** GFP to mRFP fluorescence ratio in EGFP-LC3B-mRFP-LC3B1′G transduced cells cultured in 2D with none, 100 nM bafilomycin A1 (baf1) or 200 nM Torin1 for 1 day or cultured in 3D with none (3D+none) or CM (3D+CM) for 7 days. Stars indicate the significance of the difference with untreated 2D-cultured (control) cells. **G)** Data from a representative flow cytometry experiment showing the distribution of the ratio of GFP to mRFP fluorescence in transduced cells cultured as indicated. **H)** Clonogenic assays to compare SSQ induction in cells pretreated at medium cell density in 2D-cultures with none or 200 nM Torin1 for 1 day (pretreatment) before cell passaging at low density. Labels indicate the percentage of seeded cells having formed a cell clone after SSQi supplementation. Stars indicate statistical significance of differences with 2D control in panel **D** and with untreated cells in panel **H** in clonogenic tests with SSQi supplementation.

### Beclin-1 expression is required for SSQ induction in an autophagy-independent manner

To further investigate the possible role of autophagy in regulating SSQ induction, we focused on Beclin-1 (BECN1) that promotes autophagosomal membrane nucleation ^17^. Beclin-1 is released from the cytoskeleton upon mTOR Complex 1 (mTORC1) inhibition and is also required for induction of autophagy by cell detachment from extracellular matrix ^14, 18, 19^. We generated 3 independent KC-DT66066 cell populations transduced with a *Becn1*-targeting lentiviral shRNA that exhibited a 90% reduction in *Becn1* mRNA level, compared to cell populations transduced with a *gfp*-targeting control lentiviral vector (KCsh*Becn1* and KCsh*gfp* cell populations, respectively; data not shown and see below). Clonogenic assays as performed in Fig 1 showed that induction of SSQ was significantly decreased by *Becn1* downregulation (Fig 3A). Furthermore, this could not be alleviated by Torin1 pre-treatment of cancer cells, which increased the frequencies of cells in SSQ only in control cell populations (KCsh*gfp*, Fig 3B). Similar results were obtained using KC-A338 cells in which a four-fold reduction in *Becn1* mRNA levels decreased SSQ induction 2.7-fold (Fig S2C).

**Fig 3:**
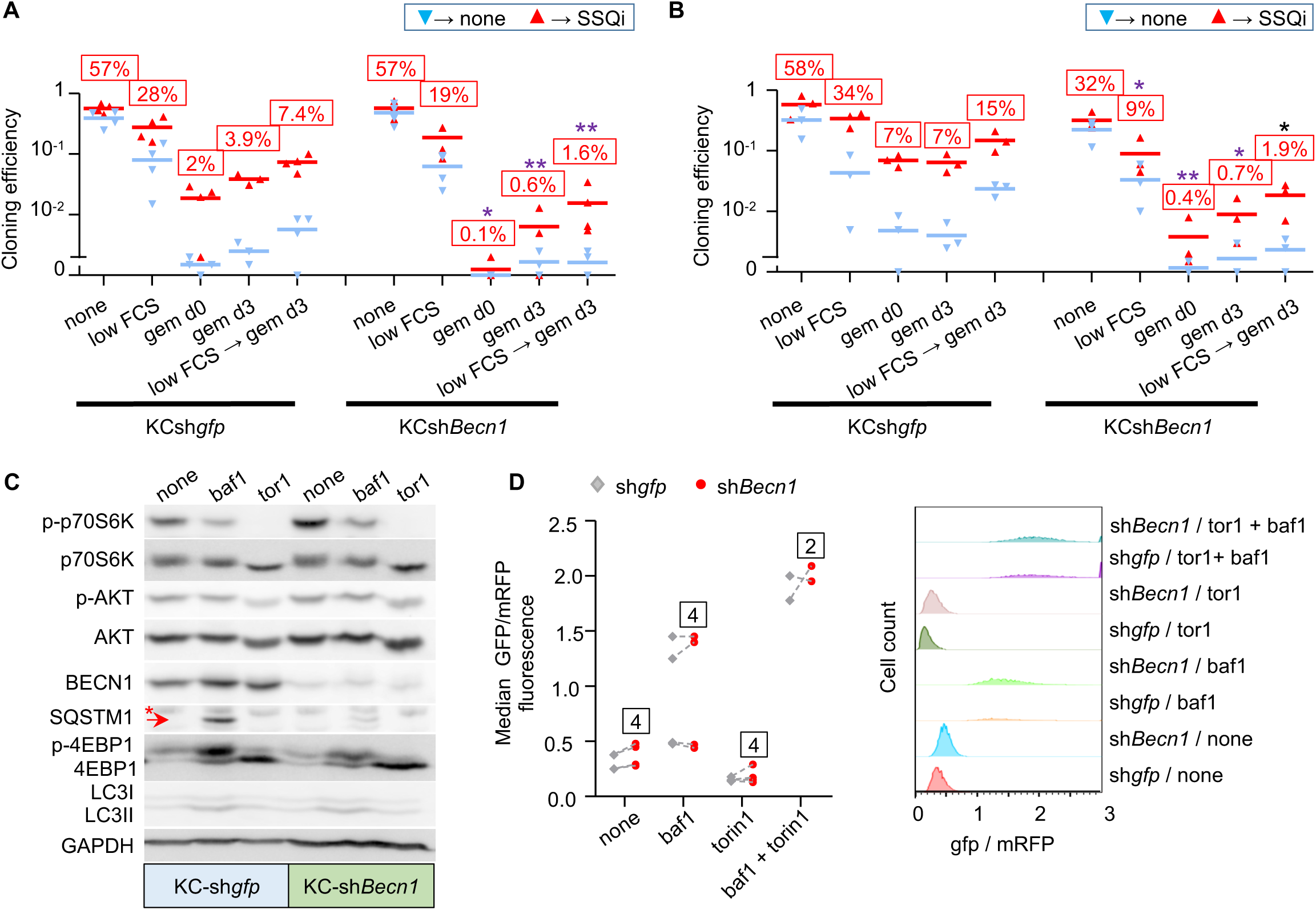
Beclin-1 downregulation inhibits SSQ but not autophagy. **A)** and **B)** clonogenic assays to compare SSQ induction in KC-DT66066 cell populations transduced with shRNA targeting GFP (KCsh*gfp*) or Beclin-1 (KCsh*becn1*) and pretreated with none **(A**) or 200 nM Torin1 (**B**) as depicted in Fig 2H. Labels above bars indicate the percentage of seeded cells having formed a cell clone after SSQi supplementation. * indicate statistical significance of CE differences after SSQi supplementation between KCsh*gfp* and KCsh*becn1* cells. **C**) Western Blot analysis of KCsh*gfp* and KCsh*Becn1* cell populations seeded at medium cell density at d0 and treated from d1 to d2 as indicated (baf1 and tor1 were for 100 nM bafilomycin A1 and 200 nM Torin1 respectively). This is one representative western blot of 3 analyzed. The arrow indicates the SQSTM1 band and the star a non-specific band. **D**) *left*: GFP/mRFP fluorescence ratio showing no difference in autophagic flux in KC*shgfp* and KC*shbecn1* cell populations transduced with the EGFP-LC3B-mRFP-LC3BΔG reporter gene. Cells were passaged at medium cell density at d0, treated from d1 to d2 with the indicated drugs (none, bafilomycin A1 (baf1), and/or Torin1) and analyzed as in Fig 2F-G. Dotted lines connect data points derived from a same experiment. Labels indicate the number of independent reporter gene transduction experiments. *Right*: A representative flow cytometry experiment shown.

Western Blot analysis showed that KCsh*Becn1* cell populations displayed a 10-fold decrease of Beclin-1 (Fig 3C). Phosphorylation of p70S6 kinase and 4EBP1, two downstream targets of mTOR complex 1 (mTORC1) was strongly inhibited by Torin1 treatment in both control and KC*shBecn1* populations (Fig 3C). Akt1 phosphorylation at serine 473 was less affected by Torin-1 treatment (Fig 3C), consistent with phosphorylation of this site by several kinases in addition of mTOR complex 2 ^20^. Interestingly, the expression of LC3-II (a cleavage product of LC3-I (ATG8) and a marker of autophagosomes formation) showed no significant difference between control and KCsh*becn1* cells under basal conditions or after treatment with bafilomycin A1 (an autophagy inhibitor that induced an accumulation of LC3-II ^21^; Fig 3C). Additionally, the level of SQSTM1, a protein degraded by autophagy, was barely detectable under basal conditions in both control and KCsh*becn1* cells, but was increased by bafilomycin A1 treatment (although to a lesser extent in KCsh*becn1* cells; Fig 3C). This suggested that autophagy was barely affected by Beclin-1 downregulation, which was confirmed by comparing autophagic flux in several independent KCsh*gfp* and KCsh*Becn1* cell populations (Fig 3D). Thus, Torin1 treatment could promote SSQ through Beclin1-dependent mechanisms that are unrelated to autophagy. This was confirmed by two types of experiments. Inhibition of autophagic flux with bafilomycin A1 (which affects autophagy downstream of Beclin-1 activation) did not have a significant impact on the induction of SSQ by Torin1 in pancreas or prostate adenocarcinoma cells (Fig S2D-E). Similarly, transduction of a constitutively active mutant of Beclin1, Becn1*^F121A^*^22^ which is defective in its binding to Bcl2/Bclx inhibitors, resulted in increased autophagic flux both under basal conditions or after mTOR inhibition, but did not have a significant impact on SSQ induction (Fig S2F-S2H).

### Cancer cells in SSQ are distinguished from senescent cells and exhibit Epithelial to Mesenchymal Transition (EMT) characteristics

Since anticancer agents are potent inducers of replicative senescence in cancer cells, we investigated a possible relationship between replicative senescence and SSQ by characterizing cancer cells in SSQ after gemcitabine selection. Thus, KC-DT66066 cells were treated for 1 day with Torin-1 at medium cell density and then individually tracked after SSQ induction at low cell density using the low FCS⟶ gem d3⟶ SSQi clonogenic assay. Cancer cells in SSQ were identified *post hoc* as the single cells at day 7 that survived gemcitabine selection and then gave rise to a cell clone after culturing in growth medium supplemented with SSQi. Accordingly, cells were photographed at the end of gemcitabine selection on day 7 (d7) and after culturing in growth medium supplemented with SSQi on days 10, 14 and 17 (Fig 4A). It appeared that most of the clonogenic cancer cells in SSQ were small spindle-shaped or rounded cells (Fig 4B upper row and table 1). Non-clonogenic cancer cells showed more variable morphology, with significantly enlarged nucleus and cell area, which are two distinctive features of senescent cells ^23–25^ (Fig. 4B lower row and Fig 4C). In fact, a large proportion of non-clonogenic cells were large, flattened senescent-like cells whereas only 1 of 29 clonogenic cancer cells in SSQ exhibited characteristics of senescent cells (Fig. 4B and Table 1), suggesting that SSQ and replicative senescence are disjoint phenomena. This was consistent with the reversible nature of SSQ (Fig S1F) whereas senescent-like cells appeared to have undergone irreversible growth arrest. Indeed, two independent cell populations highly enriched in cells with a senescent phenotype by culturing them for 10 days in the presence of gemcitabine at medium cell density showed a strong decrease below the detection threshold in their cloning efficiency and their ability to enter SSQ.

**Fig 4:**
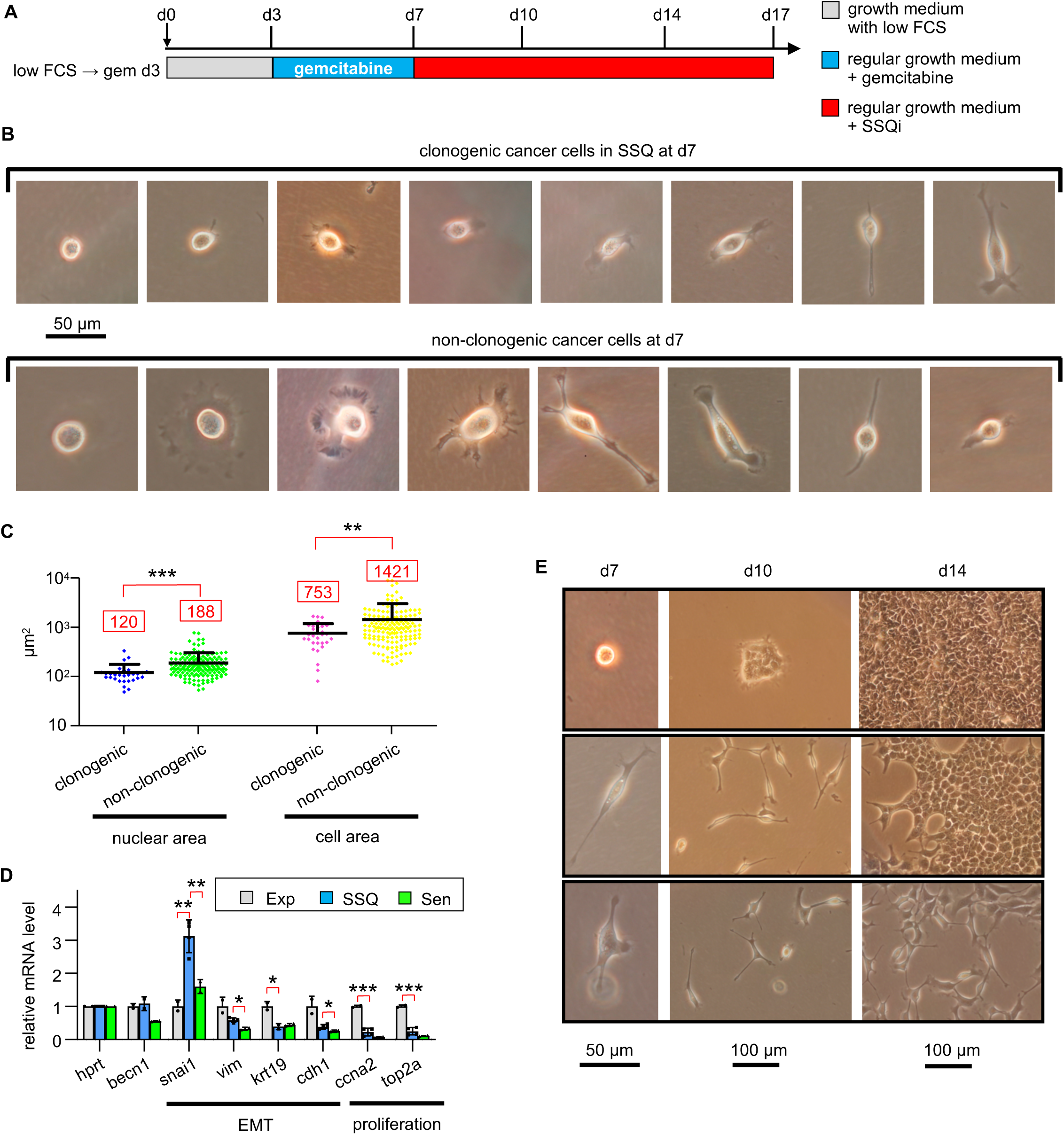
KC-DT66066 cells in SSQ are distinguished from senescent cells and exhibit morphological characteristics of Epithelial to Mesenchymal Transition (EMT). **A)** Outlines of cancer cell tracking in the low FCS⟶ gem d3⟶SSQi clonogenic assay using a phase contrast microscope to identify cancer cells in SSQ. Cells were pretreated for 1 day with 200 nM Torin1 at medium cell density before seeding at low cell density and in low serum at day 0 (d0). The cancer cells in SSQ were identified *a posteriori* as individual cells at the end of gemcitabine selection (at d7) having given rise to a cell clone after culture in growth medium supplemented with SSQi. **B)** Representative pictures of clonogenic cancer cells in SSQ (upper row) and non-clonogenic cancer cells (lower row) photographed on day 7. Scatter plots showing greater nuclear and cellular size in non-clonogenic cells at day 7 compared to SSQ cells. A total of 186 cells were analyzed in two independent experiments. RT-qPCR analysis of mRNA species levels in SSQ-enriched cell populations (SSQ; n=4) versus growing cell populations (Exp; n=2) and senescent cell populations (Sen; n=2). SSQ-enriched cell populations were Torin1-pretreated cells from the low FCS⟶ gem d3 clonogenic assay analyzed at day 7. Senescent cells populations were derived by culturing cells at medium cell density with gemcitabine plus SSQi for about 10 days and further culturing for 1 day in regular medium before cell lysis. Stars indicate statistical significance of the differences with SSQ-enriched cell populations using a two tailed heteroscedastic student’s t test. E) Representative pictures of cancer cells in SSQ on day 7 that have transitioned to an epithelial phenotype by day 10 (upper row), day 14 (middle row) or have maintained a mesenchymal phenotype by day 14 (lower pictures row).

**Table 1:**
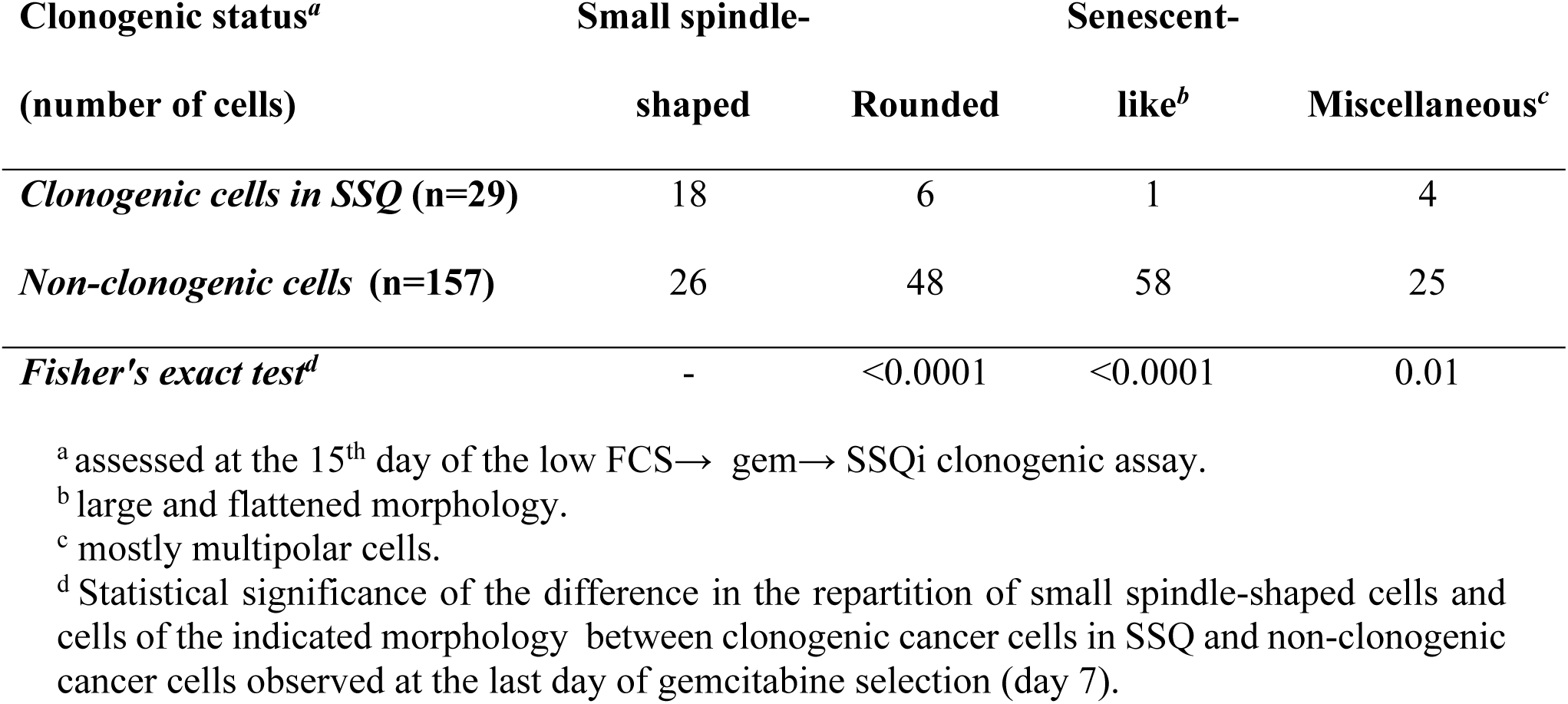
Most clonogenic KC-DT66066 cells in SSQ are small spindle-shaped cells.

| **Clonogenic status<sup>a</sup>**<br>**(number of cells)** | **Small spindle-**<br>**shaped** | **Rounded** | **Senescent-**<br>**like<sup>b</sup>** | **Miscellaneous<sup>c</sup>** |
| --- | --- | --- | --- | --- |
| *Clonogenic cells in SSQ (n=29)* | 18 | 6 | 1 | 4 |
| *Non-clonogenic cells (n=157)* | 26 | 48 | 58 | 25 |
| *Fisher's exact test<sup>d</sup>* | - | <0.0001 | <0.0001 | 0.01 |

<sup>a</sup> assessed at the 15<sup>th</sup> day of the low FCS→ gem→ SSQi clonogenic assay.
<sup>b</sup> large and flattened morphology.
<sup>c</sup> mostly multipolar cells.
<sup>d</sup> Statistical significance of the difference in the repartition of small spindle-shaped cells and cells of the indicated morphology between clonogenic cancer cells in SSQ and non-clonogenic cancer cells observed at the last day of gemcitabine selection (day 7).

The spindle-shaped of most cancer cells in SSQ suggested that these cells had undergone an Epithelial-Mesenchymal Transition. This was consistent with increased expression of *Snai1*, a potent EMT inducer, and decreased expression of epithelial markers (*Cdh1 and Krt19*) in SSQ-enriched cell populations compared to proliferating cells (Fig. 4D). Senescent cells populations derived from medium cell density cultures treated with gemcitabine plus SSQi for about 10 days showed no significant *Snai1* induction highlighting that *Snai1* induction in SSQ-enriched cell populations did not result of their contamination by senescent cells (Fig 4D). The prevalence of the mesenchymal phenotype in SSQ cancer cells was supported by the observation that 26 of 29 growing cell clones that formed after 3 days of culture in SSQi-supplemented medium (on day 10; n= 13±7.3 cells/clone; median ± standard deviation) were composed of scattered spindle-shaped cells whereas only 3 clones were epithelial type (see for example Fig 4E, middle column). The persistence of a mesenchymal phenotype over multiple cell generations was remarkable since native KC-DT66066 cultured at medium cell density exhibited a marked epithelial phenotype (see for instance Fig S1C). However, 25 of 29 cell clones had switched to an epithelial phenotype by day 14 and 28 of 29 by day 17 (see also Fig 4E, right column). Interestingly, a delayed transition to an epithelial phenotype was significantly associated to a low clonal cell growth rate, since the 4 clones that transitioned after day 14 were small compared to only 8 of the 24 that transitioned before day 14 (p<0.02 using a Fisher’s exact probability test; see Fig 4E for an instance).

### SSQ regulates tumor growth in vivo in mice

When 10^6^ carefully dissociated KC-DT66066 cells harvested from growing cultures were subcutaneously injected in allogenic C57Bl/6 mice, they gave rise to fast-growing tumors that began to regress after approximately one week due to immune rejection of cancer cells. Even during this short period of growth, we observed a higher tumor growth rate with Beclin-1 deficient cell population (KCsh*Becn1*) compared to the control (KCsh*gfp*) (Fig S3A). Interestingly, we found that treatment with a high dose of gemcitabine on day 4 after cell injection increased KC cell tumorigenicity, possibly by depressing the immune response (data not shown and see ^26^). Under these conditions, the higher growth rate of KCsh*Becn1* cells was confirmed because KCsh*becn1* cells, but not the control population, were able to out-compete the immune response (Fig S3B). Considering that KCsh*becn1* and KCsh*gfp* cell populations exhibited the same growth rate in 2D culture at medium cell density (Fig S3C) and the same percentage of clonogenic cells (compare none-+ SSQi in Figs 3A and 3B), these experiments suggested that a significant fraction of control cells could enter SSQ *in vivo*, resulting in a lower tumor growth rate. Of note, they also showed that KC-DT66066 cells exhibit high resistance to gemcitabine *in vivo*, which is not dependent on Beclin-1 expression. To further investigate the role of SSQ in regulating tumor growth, we analyzed the tumorigenicity of KC-DT66066 cells in immunodeficient NOD-SCID mice. As few as 100 dissociated cells harvested from exponentially growing cell cultures gave rise to rapidly growing tumors in these hosts (Fig S3D). To test the contribution of cells in SSQ, we subcutaneously injected 4 mice with 500 cells obtained from exponentially-growing culture (which were refer to as the Exp group that additionally included 2 mice previously injected with 100 proliferating cells, n=6) and 8 mice with 500 cells enriched for SSQ by Torin 1 treatment and gemcitabine selection in cell culture (Fig 5A and Methods). Mice injected with cells enriched for SSQ were further divided into 2 groups that were treated with saline (Ssq/none group, n=4) and gemcitabine on the 4^th^ and 11^th^ day after cell injection (Ssq/gem group, n=4; Fig 5A and Methods). Gemcitabine treatment was performed with the aim of counter selecting cancer cells that started to proliferate shortly after cell injection to further enrich the injected cells for SSQ cells *in vivo*. A distinct palpable tumor mass (≈10 mm^3^) could be detected in all the mice in the Exp and Ssq/none groups and in 3 of the 4 mice in the Ssq/gem group around the 30^th^ day after cell injection (Fig 5B-C). These results indicated that the initial phase of tumor growth was not significantly affected by the state of the cells prior to injection or by treatment with gemcitabine. However, subsequent tumor growth differed in the groups of mice injected with SSQ cells. While all tumors in mice of the Exp group grew vigorously and rapidly reached a tumor volume of 100 mm^3^ approximately one week later (36.7 ± 3.9 days, mean ± s.d., n=6), 2 tumors out of 4 in mice in the SSQ/none group thereafter grew very slowly and reached 100 mm^3^ only at day 67 after cell injection. This corresponded to a difference with the mean tumor growth time of the Exp group of more than 7 times the standard deviation of the tumor growth times in the Exp group (Fig 5B and 5D). Such a difference makes the occurrence of slow-growing tumors unlikely in the Exp group if a Gaussian distribution of tumor growth rates in this group is assumed. Additionally, a third mouse in the Ssq/none group developed a tumor that rapidly reached 200 mm^3^ on day 37, but then stopped growing until this mouse was sacrificed on day 47 due to skin ulceration (Fig 5B). The proportions of slow-growing tumors in the Exp and Ssq/none groups (0/6 *vs* 3/4) were found statistically different as calculated with a Fisher’s exact probability test (p=0.033). This confirmed that formation of tumors with a slow-growing phenotype is promoted by the injection of cancer cells in SSQ. Similar observations were made with mice in the Ssq/gem group (3 of 4 mice also developed slow-growing tumors), but one mouse also showed delayed formation of a palpable tumor (it was detected on day 65 after cell inoculation; Fig 5B) in addition to slow tumor growth (Fig 5D). This showed that cancer cells in SSQ promoted the formation of indolent tumors at high frequency and that SSQ could also impede the early phase of tumor growth. Nonetheless, all of the indolent tumors, including the one that had been indolent for two months, eventually transitioned to a rapidly growing phase, consistent with our concept of a stable but reversible quiescence (Fig 5B). Genes expression profiling of growing tumors by RT-qPCR analysis did not reveal any significant difference associated with the status of the injected cells (correlation coefficient r>0.99, n = 17 genes analyzed, p <0.0001; Fig S3E and S1 file). Interestingly, in comparison with exponentially growing KC-DT66066 cells in 2D-cultures, all growing tumors displayed an increased expression level of the *Lgr5* stemness marker, as observed for cells growing in spheroids (compare Figs 2C and S3E). This analogy between spheroids and tumors was further supported by analysis of the clonogenicity of dissociated tumor cells, as previously done for spheroids (Fig 2D, 5E and see Methods). This not only confirmed the presence within the tumors of significant populations of cells in SSQ (18.7± 8.2% of clonogenic tumor cells selected with the gem d0 clonogenic assay, mean ± sd, n=5; Fig 5E), but also revealed that clonogenic tumor cells were prone to enter SSQ (27.3 ± 11.4% of clonogenic tumor cells selected with the low FCS→ gem d3→ SSQi clonogenic assay, mean ± sd; n=5; Fig 5E). Interestingly, in tumors that grew in allogenic C57Bl/6 mice injected with 10^6^ proliferating KC-DT66066 cells, clonogenic assays also revealed cancer cells in SSQ or prone to enter SSQ after seeding at low cell density under low serum, but at lower frequencies (1.3 ± 0.7% and 3.0 ± 1.7% of clonogenic cells, respectively, mean ± sd, n=5). However, these frequencies significantly increased when C57Bl/6 mice were treated with cyclosporine, an immunosuppressive agent (4.50±1.1% and 14.8±4.9% of clonogenic cells, respectively; n=4, p<0.002; see Methods for more details), suggesting that immune response could modulate frequency of SSQ cells in tumors.

**Fig 5:**
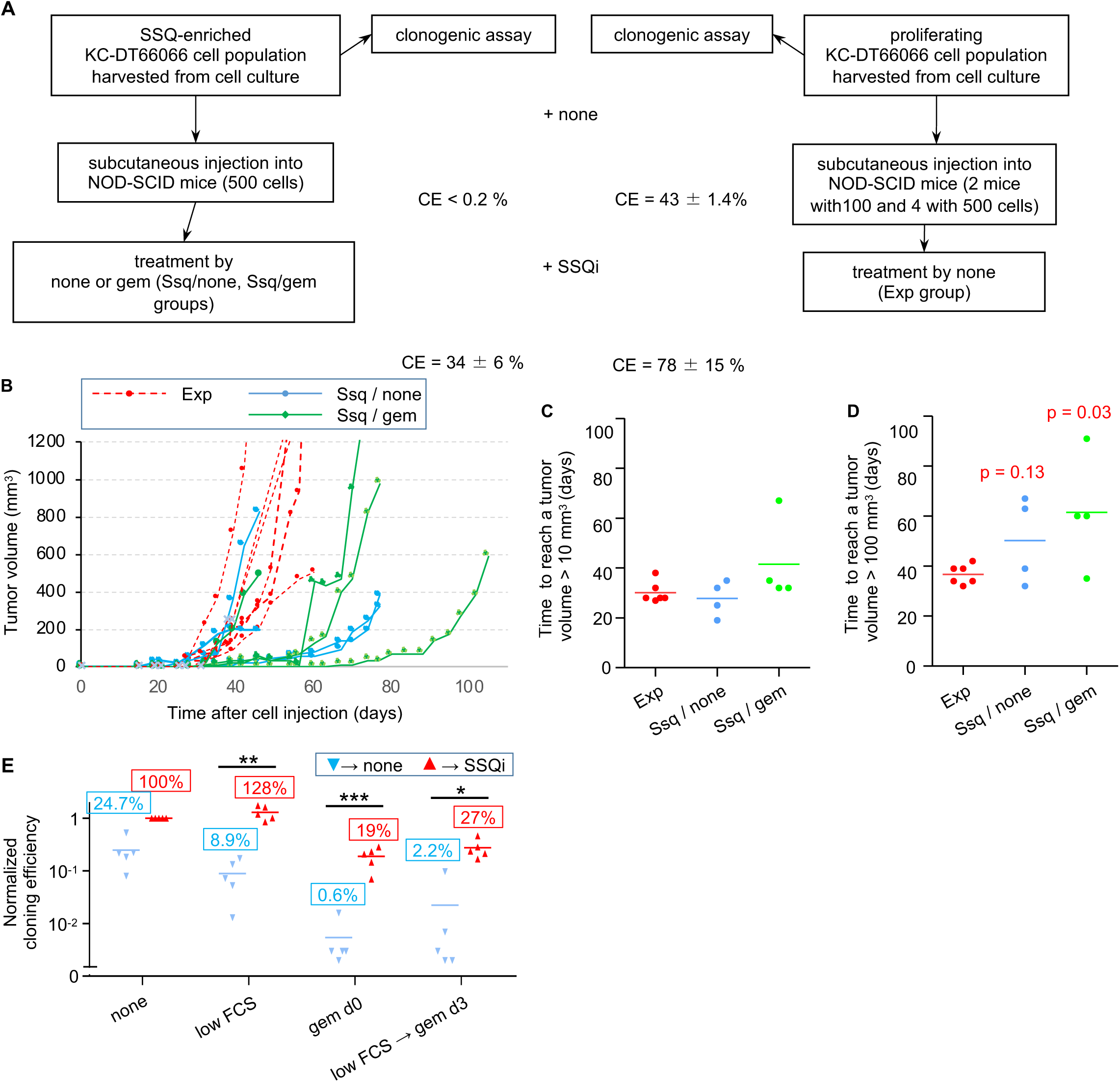
SSQ regulates tumor development. **A)** Outlines of the experiment performed to compare tumorigenicity of KC-DT66066 cells in a proliferating state or in SSQ. **B)** Tumor growth curves in individual NOD-SCID mice subcutaneously injected with proliferating KC-DT66066 cells (Exp group, n=6) or with cells in SSQ and treated with none (Ssq / none group, n=4) or gemcitabine (Ssq / gem group; n=4). **C**) and **D**) Scatter plots showing the time taken for tumors to reach a volume of 10 mm^3^ (**C**) or 100 mm^3^ (**D**). p indicates the statistical significance of the difference in the proportion of mice with a slow tumor growth rate as calculated with a Fisher’s exact probability test. Note that a third tumor in the Ssq/none group showed a slow tumor growth rate but only after having reached 200 mm^3^ (see Fig 5B and main text). **E**) Growing tumors contain a significant percentage of cancer cells in SSQ or prone to enter SSQ. Clonogenic assays using cells dissociated from growing tumor resected from mice in the Exp (n=1), Ssq/none (n=2) and Ssq/gem (n=2) groups and directly tested for SSQ as described in Fig 1A. Data were normalized to 100% of cloning efficiency in the none ⟶ SSQi clonogenic assay. * indicates the statistical significance of the change in CE induced by SSQi supplementation. Note that spontaneous reversion of SSQ (that could be observed with the clonogenic assay low FCS → gem d3→ none) strongly increased in some dishes containing a high absolute number of cells in SSQ as observed in Fig 2D.

### High Beclin-1 level and low mTOR activity in the primary tumor of PAAD patients is predictive of late metastatic recurrence

Based on our results in mice, primary tumors with a relatively low level of Beclin-1 expression should release cancer cells that are not prone to enter SSQ, and thus cause a lower rate of late metastasis if SSQ constitutes the only mechanism controlling delayed metastases development in pancreatic adenocarcinoma (PAAD). Analysis of PAAD patient records downloaded from the Genomic Data Commons website (see Methods) showed that the rate of distant metastases in the first 360 days after initial treatment did not differ significantly between cohorts of patients with low and high level of Beclin-1 transcripts in their primary tumors (Fig 6A). However, beyond 360 days, a low level of Beclin-1 transcripts in the primary tumor was associated with a decreased rate of distant metastases (Fig 6B and see the Methods section for the determination of the cutoff value of *BECN1* mRNA levels used throughout this study). This remained true when we restricted the analysis to patients with a similar T3N1M0/Mx clinical grade (Fig S4A-B), suggesting that Beclin-1 expression level is a predictive factor of late metastasis independent of TNM grade. To further establish the biological relevance of a 360-day cutoff for classifying patients as developing early or late metastases, we compared the primary tumors transcriptomes of patients with distant metastasis-free survival less than and greater than 360 days by Gene Set Enrichment Analysis (GSEA). GSEA recovered only 8 statistically significant genes sets with a false discovery rate (FDR) q<0.05. They showed that primary tumors from patients with a distant-metastasis-free survival inferior to 360 days were enriched for genes regulated by MYC transcription factor, mTORC1 signaling and the unfolded protein response (Fig 6C and see Fig S4C for others miscellaneous gene sets). This suggested that a distinguishing feature of tumors from patients with a distant metastasis-free survival inferior to 360 days might be enrichment in proliferating cells (enhanced MYC activity) that exhibited higher protein synthesis (enhanced mTORC1 signaling) associated with endoreticulum stress (enhanced unfolded protein response). Thus, we analyzed distant metastasis-free survival of PAAD patients according to the relative level of Cyclin A2 (*CCNA2*, a marker of cell proliferation) in their primary tumor since mTORC1 activity seemed to be associated to tumor cell proliferation in the primary tumors. A high level of Cyclin A2 transcript was found to be a very strong predictor of early metastasis with a median survival time of 291 days *versus* 763 days in the cohort of patients with a low level of Cyclin A2 transcript (Fig 7A and see the Methods section for the determination of the cutoff value of CCNA2 mRNA levels used throughout this study) and this was independent of TNM grade (Fig S4D). Independence was also confirmed by multivariate Cox regression analysis (data not shown). GSEA showed that primary tumors with a high level of Cyclin A2 transcripts were highly enriched for classical hallmarks of cell proliferation, but also enhanced mTORC1 signaling and unfolded protein response (Fig 7B and S4E), as anticipated above. The strong correlation between the transcript levels of Cyclin A2 and of markers of mTORC1 activity in the primary tumors of PAAD patients were confirmed by hierarchical gene clustering (Fig 7C). Interestingly, this analysis also showed that levels of Beclin-1 and Cyclin A2 transcripts (as well as mTORC1 activity marker genes) were positively correlated with levels of cancer cell markers but not mesenchymal, endothelial or immune cell markers (Figs 6C and S4F). This suggested that transcript levels of Beclin-1 and Cyclin A2 in the primary tumors were mainly determined by cancer cells.

**Fig 6:**
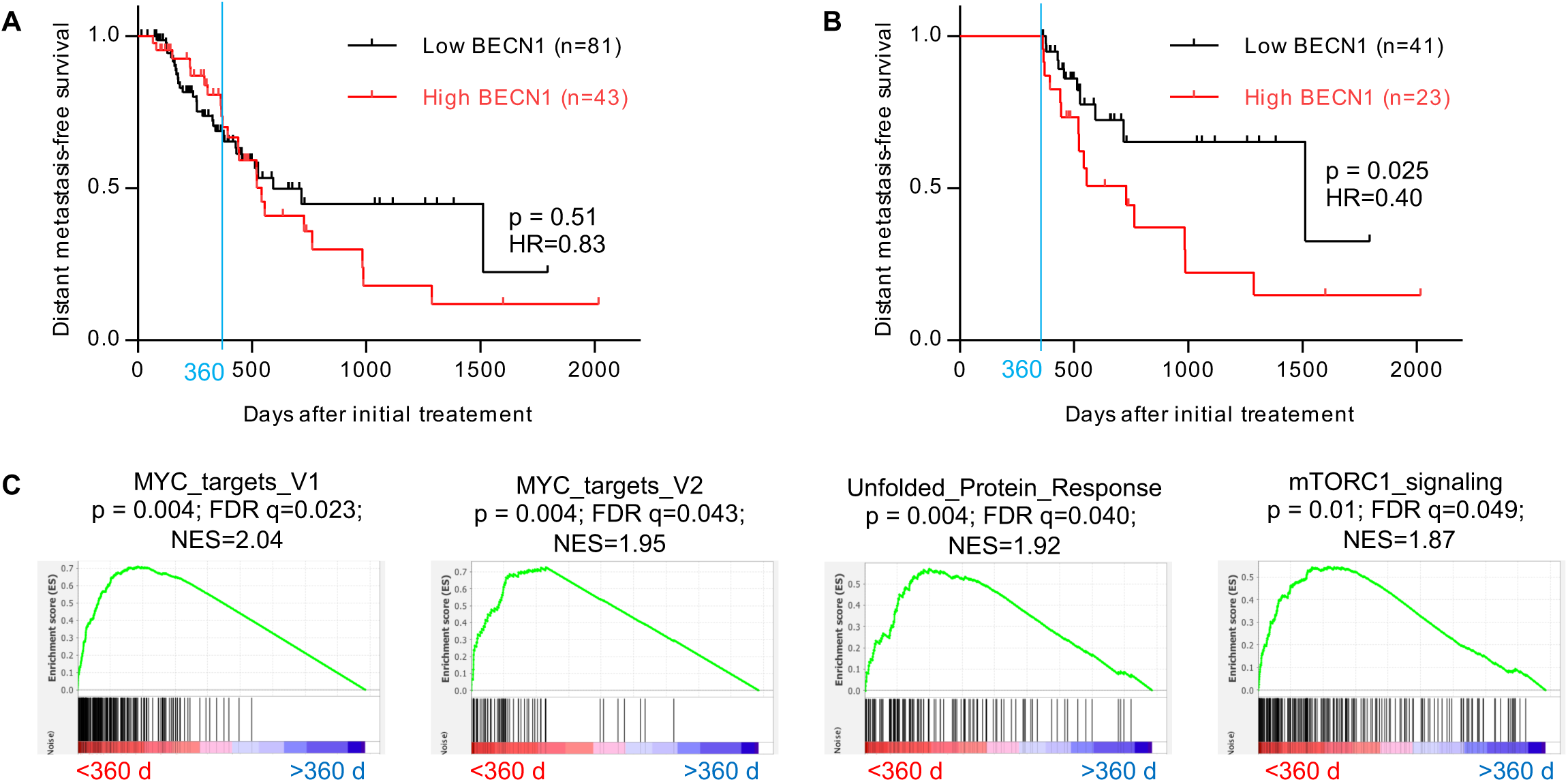
Elevated Beclin-1 expression in the primary tumor of PAAD patients is associated with an increased rate of metastasis in the patient cohort with a metastasis-free survival greater than 360 days. **A)** Kaplan-Meier curves for distant metastasis-free survival in PAAD patients according to the level of Beclin-1 transcripts in their primary tumor. **B)** Same as **(A)** but restricted to patients with survival time greater than 360 days. Initial treatment was partial or total pancreatectomy to remove the tumor and at least 78% of patients received adjuvant pharmaceutical treatments (see **S2 file**). Significance (p) of the difference in survival and hazard ratio (HR) were calculated with a Mantel-Cox Log-ranked test. **C)** Gene set enrichment analysis (GSEA) showing significant enrichment for hallmarks of MYC, mTORC1 and unfolded protein response signaling activity ^57^ in primary tumors of PAAD patients with a distant-metastasis-free survival time less than 360 days after initial treatment (n=60) *versus* greater than 360 days (n=64). Normalized enrichment score (NES), its statistical significance (p) and its false discovery rate (FDR) are indicated.

**Fig 7:**
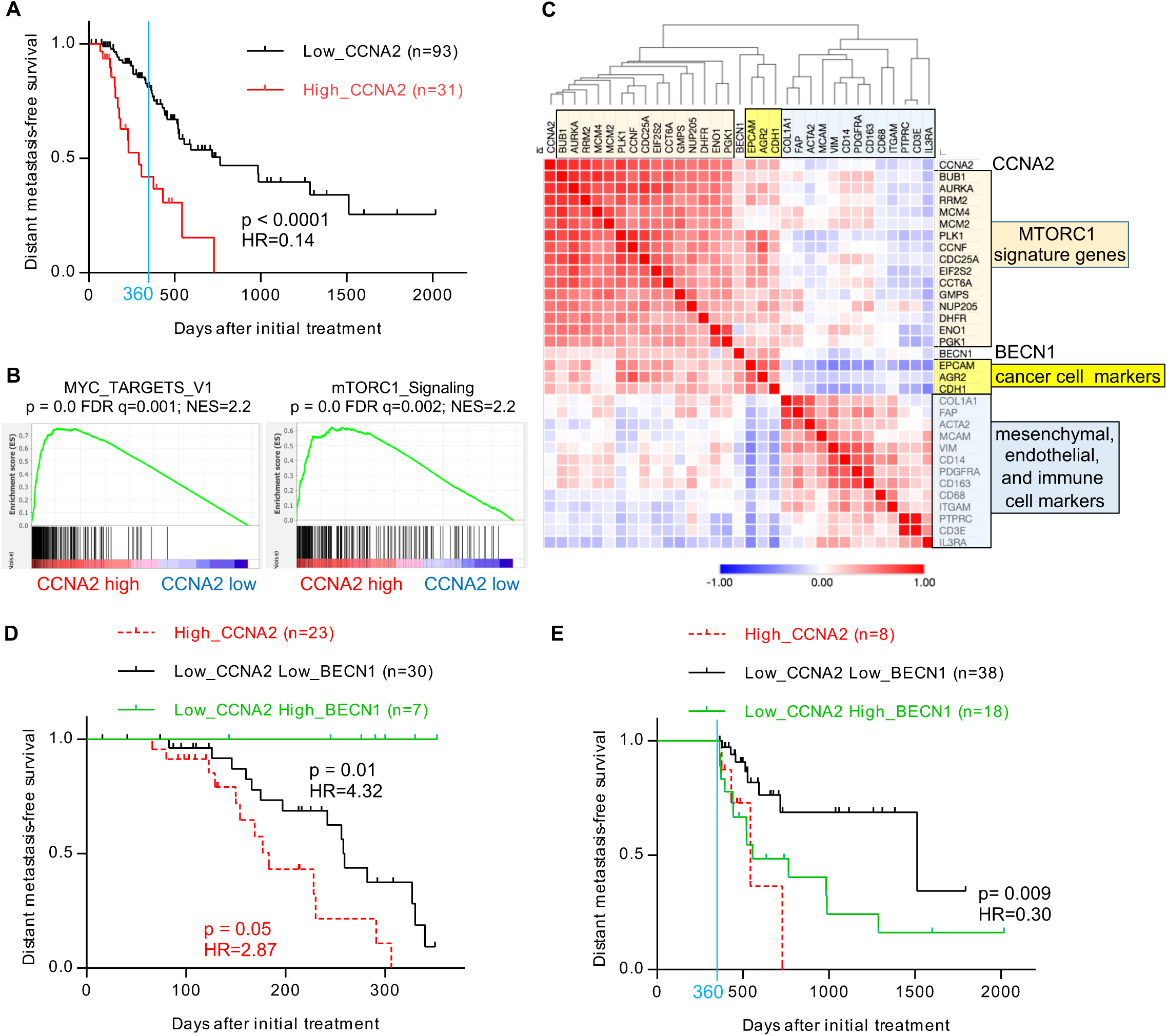
Cyclin A2 marks tumors with elevated mTORC1 activity and combines with Beclin-1 to predict kinetics of metastases development. **A)** Kaplan-Meier curves for distant metastasis-free survival in PAAD patients with high or low level of Cyclin A2 (*CCNA2*) transcripts in their primary tumor. **B)** GSEA demonstrates significant enrichment for hallmarks of cell proliferation and mTORC1 signaling ^57^ in the primary tumors of PAAD patients with high level (n=31) versus low level (n=93) of *CCNA2* transcripts. Normalized enrichment score (NES), its statistical significance (p) and its false discovery rate (FDR) are indicated. **C)** Hierarchical clustering of genes according to the Pearson correlation of their transcription levels in the primary tumors of PAAD patients (n=124). The colors indicate the value of the Pearson correlation coefficient for a given gene pair, ranging from −1 (blue) to +1 (red). The mTORC1 genes markers correspond to the first 15 highest ranked genes composing the mTORC1 signaling signature derived by GSEA analysis of PAAD tumors in Fig 6C (gene set M5924; ^57^). **D)** Kaplan-Meier curves for distant metastasis-free survival in PAAD patients with survival time less than 360 days according to the transcript levels of cyclin A2 (CCNA2) and Beclin 1 (BCEN1) in their primary tumor. **E)** Same as panel D), but limited to patients with a survival time greater than 360 days. Log-ranked (Mantel-Cox) test was used to calculate the statistical significance (p) of the differences in distant metastasis-free survival between adjacent survival curves and hazard ratios (HR).

The combination of Beclin-1 and Cyclin A2 markers showed that all metastatic events within the first 360 days after initial treatment were associated with tumors with either high Cyclin A2 or both low Cyclin A2 and low Beclin-1 expression (Fig 7D). Remarkably, patients whose tumors exhibited both low Cyclin A2 and high Beclin-1 expression did not develop metastases within the first 360 days after initial treatment (Fig 7D). However, after 360 days, patients with both low Cyclin A2 and high Beclin-1 expression exhibited a higher rate of distant metastasis compared to patients with both low Cyclin A2 and low Beclin-1 expression in their primary tumors (Fig 7E). Taken together, this strongly suggests that Beclin-1 abundance and mTORC1 activity in the primary tumor of PAAD patients determine the kinetics of occurrence of clinical metastases. This was further confirmed by analyzing the statistics of the relative proportions of early and late clinical metastasis (before and after 360 days) according to tumor phenotype of patients with a T3N1M0/x clinical grade (Table 2). This showed that late clinical metastasis was mainly associated to the *CCNA2^low^ BECN1^high^* primary tumor phenotype and early clinical metastasis to the *CCNA2^high^*and the *CCNA2l^ow^ BECN1^low^* primary tumor phenotypes.

**Table 2:** Early and late (before and after 360 days, respectively) distant metastases events recorded in T3N1M0/x PAAD patients according to their primary tumor phenotype.

| primary tumor phenotype<br>(number of patients) | SSQ<br>competence | Early metastases<br>events<sup>a</sup> | Late metastases<br>events<sup>b</sup> | Fisher's<br>exact test<sup>c</sup> |
| --- | --- | --- | --- | --- |
| *CCNA2*<sup>high</sup> (n=20) | low | 8 | 2 | 0.0007 |
| *CCNA2*<sup>low</sup> *BECN1*<sup>low</sup> (n=41) | low | 10 | 4 | 0.0016 |
| *CCNA2*<sup>low</sup> *BECN1*<sup>high</sup> (n=18) | high | 0 | 9 | - |

<sup>a</sup> within 360 days of initial treatment.
<sup>b</sup> beyond 360 days after initial treatment
<sup>c</sup> Statistical significance of the difference in the repartition of early and late metastases events compared to the cohort of patients with a *CCNA2*<sup>low</sup> *BECN1*<sup>high</sup> tumor phenotype.

## Discussion

We showed here that cancer cells in SSQ can be selected with anticancer agents including gemcitabine, vincristine, nocodazole, doxorubicin, oxaliplatin and taxol *in vitro* (Figs 1, S1D-F) and possibly also *in vivo* (Fig 5A-B). This suggests that SSQ could promote the persistence of disseminated cancer cells after adjuvant chemotherapies. Analysis of the mechanisms of SSQ induction showed that culture at low cell density induces SSQ in prostatic and pancreatic adenocarcinoma cells (Fig 1B and ^9, 10^). By investigating why SSQ is also induced at high cell density in non-growing cell aggregates or in growing cancer cell spheroids generated in 3D culture (Fig 2D), we found that it is induced by mTOR inhibition. Indeed, pretreatment of growing cancer cells cultured in 2D at medium cell density with rapamycin or Torin1, two standard pharmacological inhibitors of mTOR activity, is sufficient to induce SSQ and also synergizes with low cell density for SSQ induction (Figs 2H and S2A-B). As cell proliferation depends on mTOR activity ^27–29^, this implies that proliferating cells are not prone to enter SSQ, in agreement with our finding that exponentially growing cell cultures in 2D exhibited low frequency of cancer cells already in SSQ and are modestly induced to enter SSQ after seeding at low cell density (Fig 1B). Inversely, in 3D cultures, high levels of autophagy indicate that mTOR activity was downregulated in a large fraction of cells since mTOR activity inhibits autophagy. An unexpected result is that induction of SSQ requires Beclin-1 expression but was independent from autophagy. Indeed, downregulation of Beclin-1 inhibits SSQ induction but not of autophagy (Fig 3) and autophagy inhibition with bafilomycin A1 does not prevent SSQ induction by Torin1 (Fig S2D-E). Moreover, transduction of a constitutively active Beclin-1 mutant, defective in its binding to Bcl2/Bclx inhibitors, increased both basal autophagy flux and its stimulation by mTOR inhibitors, but had no effect on SSQ induction (Fig S2F-H). This confirmed that autophagic flux does not regulates SSQ induction and thus that Beclin-1 is likely to be required for SSQ through autophagy-independent pathways. Several publications have already reported autophagy-independent roles for Beclin-1. Beclin-1 regulates degradation of cell membrane receptors for growth factors and transferrin through the control of endolysosomal trafficking ^17, 30–32^. It can also inhibit the System Xc^-^ cystine/glutamate antiporter involved in cystine uptake, thus impacting cellular thiol redox potential ^33^, and also the Stat3 signaling pathway ^34^. These actions of Beclin-1 could contribute to two main features of cancer cells in SSQ which are the absence of proliferating response to stimulation by external growth factors and a decrease in the redox potential of cellular thiols. Considering that mTOR activity is regulated by nutrient depletion, cellular thiol redox potential and growth factor signaling ^35–45^, activation of Beclin-1 through inhibition of mTOR activity could therefore promote the establishment of a self-sustained inhibitory loop maintaining a low mTOR activity (Fig 8). As mTOR also directly control cell proliferation through its regulation of c-Myc activity and Cyclin D accumulation ^27–29^, our data point towards a central role of mTOR downregulation in the induction and maintenance of self-sustained quiescence (Fig 8). An issue of this study concerns the role of EMT in SSQ. We observed that pancreas cancer cells in SSQ exhibit a mesenchymal phenotype (Fig 4B, D and E). This mesenchymal phenotype is obviously not sufficient to maintain SSQ since it often persisted for several cell generations after the transition from SSQ to a proliferating state. However, it could promote the transition between proliferating and SSQ states. This would be consistent with the high frequency of the mesenchymal phenotype in SSQ cancer cells. Additionally, the slower growth of cell clones with a mesenchymal phenotype could also result from a higher rate of mesenchymal cell reentry in SSQ (see Fig 4E and the corresponding Results section).

**Fig 8:**
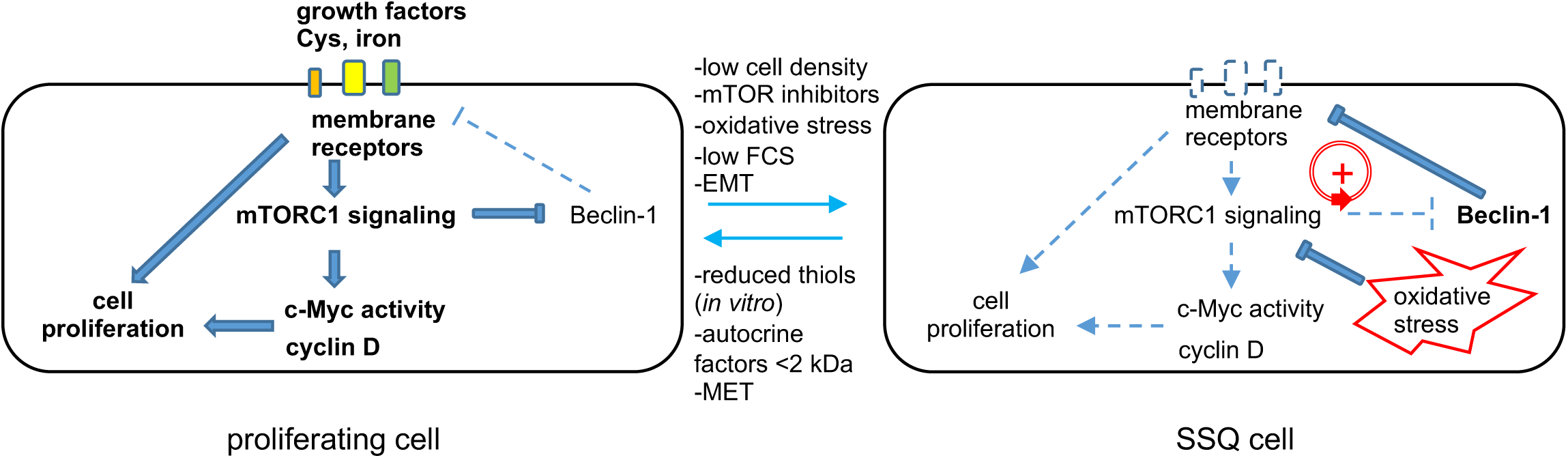
A molecular model of self-sustained quiescence. mTORC1 is a central node in the regulatory network controlling cell proliferation, for instance by regulating c-Myc activity, cyclin D accumulation and protein synthesis. Activation of Beclin-1 by downregulation of mTORC1 activity leads to activation of a regulatory loop (red circle in right panel) that helps to maintain a low mTORC1 activity. Indeed, Beclin-1 induced the degradation of cell surface receptor for growth factors and iron (transferrin receptor), and directly inhibits the Xc^-^ cystine/glutamate antiporter ^17, 30–33^, which inhibits mTOR activity via nutrient depletion, oxidative stress and suppression of growth factors signaling pathways ^35, 38, 40–45^. Note that BMP signaling stabilizes SSQ in prostate cancer cells ^8,10^ but appears to play a minor role in PAAD cells (see Methods for details and data not shown). EMT possibly favors the entry in SSQ and MET the maintenance of the proliferating state. The bold corresponds to activated signaling pathways while the dotted line corresponds to repressed pathways

Another result of our work is that dissociated PADC cells in SSQ injected into mice promoted the delayed formation of a palpable tumor in one case, as expected for the dormant phenotype of cancer cells in SSQ *in vitro*, but also, quite unexpectedly, the formation of small, slow-growing (indolent) tumors in most cases (Fig 5). Our data suggest that SSQ can directly regulate tumor cell growth and thus tumor indolence when tumor is composed of a high percentage of SSQ-competent cells. Indeed, we observed that the mouse tumor microenvironment promotes SSQ. A large fraction of clonogenic cancer cells dissociated from resected tumors were found to be in SSQ or prone to enter SSQ after cell plating at low cell density, this even in tumors resulting from the injection of exponentially growing cancer cells exhibiting a low frequency of cells in SSQ or competent for SSQ (see the Result section and also compare data in Figs 1 and 5E). This SSQ-promoting effect of tumor microenvironment could explain why tumors generated by the injection of a large number of exponentially growing cancer cells rapidly exhibit a higher growth rate when these cells have been made defective for SSQ by Beclin-1 downregulation whereas downregulation of Beclin-1 has no effect on proliferation of cells cultured at medium cell density (Fig S3A-C). Accordingly, a plausible mechanism of long-term tumor indolence is the maintenance of a high percentage of cancer cells in SSQ through the continuous induction of SSQ by tumor microenvironment (see also above the discussion on the possible role of EMT in SSQ reentry). Breaking this steady state through the rapid outgrowth of cancer cells refractory to SSQ induction or changes in the tumor microenvironment would resume a high tumor growth rate. Taken together, these results challenges models of tumor dormancy that posit a dynamic balance between cancer cell proliferation and cell death caused by antitumor immunity and insufficient tumor vascularization ^46^.

Our analysis of the relationship between metastasis in PAAD patients and gene expression in their primary tumor showed that tumors that are expected to contain and therefore disseminate clonogenic cancer cells that are not prone to enter SSQ mainly give rise to early clinical metastasis whereas those that are likely to disseminate cancer cells in SSQ or poised to enter SSQ give rise to late clinical metastasis (Figs 5-6 and Table 2). Indeed, tumors expressing a higher level of Cyclin A2, a marker of cell proliferation, also exhibit higher mTORC1 activity (Fig 7B-C), which inhibits induction of SSQ (see discussion above and Fig 8). Similarly, a low level of Beclin-1 transcripts was found to decrease the ability to enter SSQ (Fig 3), even in growth-arrested cells cultured in 3D (data not shown). Conversely, a high level of Beclin-1 expression, when combined with low mTORC1 activity associated with low Cyclin A2 expression, is expected to promote entry in SSQ (Fig 3). Of note, Cyclin A2 mRNA, in addition to being a marker of cell proliferation, is also a hallmark of c-Myc activity (it is the highest ranked gene in the GSEA analyses showing enrichment for c-Myc targets in Fig 7B and the eighteenth ranked gene in Fig 6C). Correlatively, one of the most distinctive features of patients with early metastasis is an enrichment of their primary tumor with cellular markers of c-Myc activity (Figs 6C and 7B). This suggests that c-Myc activity *per se* could be a better hallmark of mTORC1 activity in primary tumors rather than cell proliferation, which is consistent with c-Myc regulation by mTORC1 signaling via Beclin-1-dependent and independent pathways ^27, 47^.

Beclin-1 is often considered a haplo-insufficient tumor suppressor gene since Beclin-1 can inhibit tumorigenesis ^48^ and an allele of this gene is frequently deleted in advanced cancers ^49^. A high Beclin-1 expression level in the primary tumor of PAAD patients was successively associated with a low rate of metastasis in one study and then with a high rate of metastasis in another ^50, 51^. This paradox could partly result from the short follow-up of patients with high Beclin-1 in the first study, since we found that high Beclin-1 expression shows a trend to promote a lower rate of early metastasis (Fig 6A). However, it should be noted that our data support the idea that Beclin-1, combined with mTORC1 activity, primarily determines the kinetics of metastases development and not the overall rate of metastasis.

A possible direct clinical application of our results is a better selection of patients eligible for early and aggressive adjuvant chemotherapy. Indeed, our results suggest that patients with a primary tumor associated to dispersion of cancer cells not entering SSQ could benefit from aggressive chemotherapy since their dispersed cancer cells are expected to be sensitive to anticancer agents. It is crucial that adjuvant chemotherapy begins as early as possible since cancer cells grown in tumor masses gained new mechanisms of resistance to anticancer agents compared to dispersed cancer cells, including the possible *de novo* generation of cancer cells in SSQ within growing metastases microenvironment. Patients with dispersed cancer cells only in SSQ (patients with a primary tumor phenotype associated to late metastasis or treated once by an aggressive adjuvant chemotherapy) are unlikely to respond well to chemotherapy since these cancer cells are resistant to anticancer agents even when they are dispersed. Therefore, the development of new agents targeting cancer cells in SSQ seems essential. Complete elimination of dispersed cancer cells in SSQ may require the identification of agents that are effective in stably reversing SSQ *in vivo* to restore cancer cell susceptibility to chemotherapy or in directly killing cancer cells in SSQ.

## Materials and Methods

### Cells

All cells were cultured under 5% CO_2_ at 37 °C in Dulbecco’s Modified Eagle’s Medium-high glucose (4.5 g.L^-1^ glucose, glutamax 1X; #D0819 Sigma Aldrich) supplemented with fetal calf serum (10% of DMEM volume; #10270-106; various lots; GIBCO, Life Technologies), pyruvate 1 mM (#CSTVAT00-0U from EUROBIO), penicillin/streptomycin 1X (#15140-122 from GIBCO), to which distilled water was added (17.6% of DMEM volume) to make it isotonic (isoDMEM-FCS). Final fetal calf serum (FCS) concentration was 7.9% in growth medium and was decreased to 1.9% for low FCS condition by dilution with growth medium without FCS. For subculturing, cells were harvested after 5 min treatment with 0.25% trypsin-EDTA (#25200-056 from Gibco) supplemented with 1 mM pyruvate (Tryp-EDTA-Pyr), dissociated by pipetting, counted using a hemocytometer if necessary and inoculated at a dilute density. KC-DT66066, KC-A338, and KPC-A219 are independent cell lines derived from pancreas ductal adenocarcinoma (PDAC) developed in genetically engineered LSL-*Kras^G12D/+^,* Pdx1-*Cre* (KC) and LSL-*Trp53^R^*^172^*^H/+^,* LSL-*Kras^G12D/+,^* Pdx1-Cre (KPC) mouse models of spontaneous PDAC, respectively ^11, 12^. All KC and KPC-A219 cells were derived in J. Guillermet laboratory under a genetic C57Bl/6 x CD1 and C57Bl/6 x CD1 x 129 genetic background, respectively, and were authenticated by PCR. Human LNCaP* and LNCaP C4-2 (RRID:CVCL_4782) prostate adenocarcinoma were characterized previously ^10^. Cells were found to be free of mycoplasma by a PCR test carried out on their supernatant.

### Reagents

anti-Beclin-1 (clone D40C5, #3495S), anti-GAPDH (clone D16H11, #5174), anti-4E-BP1 (clone53H11, #9644), anti-phospho-4E-BP1 (Thr37/46; clone236B4, #2855), anti p70 S6 kinase (clone 49D7, #2708), anti-phospho-p70 S6 kinase (Thr389) (clone 108D2, #9234) and anti-LC3A/B (clone D3U4C, #12741) were rabbit monoclonal antibodies from Cell Signaling Technology. Anti-Akt (#9272) and anti-phospho-Akt (S473; #9271) were rabbit polyclonal antibodies from Cell Signaling Technology. Anti-SQSTM1 (#GT239) was a mouse monoclonal antibody from Invitrogen. Secondary HRP-linked anti mouse IgG (7#076S) and anti rabbit IgG (#7074S) were from Cell Signaling Technology. K02288 (#AB-M2789-10MG), Torin1 (#F6100-UBP) and rapamycin (#AB-M1768-25MG) from AbMole BioScience (USA), N-acetyl-L-cysteine (#A7250) from Sigma-Aldrich, Nocodazole (#F6120-UBP) was from UPBIO. Gemcitabine chlorhydrate and Cyclosporin A were from Sandoz (France) and Vincristine Sulfate from Hospira (France). Lipofectamine 3000 kit (#L3000-00) was from Invitrogen. LightCycler 480 SYBR Green I Master kit (#04707516001) was from Roche Diagnostics, USA)

### Clonogenic assays

2D-cultured murine pancreas and human prostate C4-2 cells were seeded at 10^3^ cells per 60 mm diameter tissue culture-treated polystyrene dish (#430169, Corning, NY, USA) for selection of SSQ cells with anticancer agents, otherwise at 10^2^ cells per dish and cultured as indicated. Human LNCaP* cells were seeded at 5000 cells per 6 cm diameter dish and cultured as indicated. To test the cloning efficiency (CE) of cells from 3D-cultures, cells were harvested, washed one time with Tryp-EDTA-Pyr and then dissociated by incubation for 20 minutes in Tryp-EDTA-Pyr at 37°C with regular pipetting. Complete growth medium was added and cells were filtered through a 70µm Cell Strainer (Falcon, Corning), washed once with complete growth medium, counted and plated at low cell density as needed. For the CE tumor cell assay, tumors were minced and cells dissociated using the same protocol as for spheroids. Only cells larger than 20 µm were counted with a hemocytometer to calculate cell dilutions for clonogenic assays (cells smaller than 20 µM were not clonogenic). In 7 of the 8 tumors analyzed, more than 90% of the clonogenic tumor cells were KC-DT66066 cells that generated easily recognizable dense and rapidly growing cell clones. In 1 tumor from the Ssq/none group that was not included in CE analysis, long-lived fibroblastic cells constituted more than 90% of the clonogenic tumor cells and generated only sparse, slow-growing fibroblastic cell clones. Cell clones were fixed in 50% ethanol and 10% acetic acid, stained with Crystal Violet and counted manually (clones with a diameter > 0.5 mm). Cloning efficiency (CE) was calculated as the ratio of the number of cell clones to that of seeded cells. Cloning efficiency in the presence of SSQ inhibitors (SSQi) was used as a measure of the percentage of clonogenic cells. The fraction of cells in SSQ in a cell population was defined as the cloning efficiency of cells from that population in the gem d0→ SSQi clonogenic assay (Fig 1A and main text). SSQi (NK, GK or GSK) consisted of a reduced thiol (N-Acetylcysteine (NAC or N) or glutathione (G) which could be used with similar efficacy at optimal concentrations) at 1 or 4 mM for murine and human cells, respectively, and of K02288 (K, an inhibitor of BMP signaling) at 100 nM. In murine PDAC cells, K02288 increased by about twofold the recovery of proliferating cell clones from cells in SSQ when NAC concentration was below 0.5 mM but was otherwise unnecessary at 1 mM NAC concentration. Similarly, SB505124 (an inhibitor of TGF7 signaling) was used with K002288 and glutathione (GSK) to reverse SSQ in LNCaP* cells, but its effect was statistically insignificant.

### 3D culture

Cells were harvested from 2D cultures by treatment with Tryp-EDTA-Pyr, counted and seeded into 24 well-plates (Cellstar, cell-repellent surface, #664970, Greiner Bio-One Gmbh) or into 6 cm dishes (Ultra-Low Attachment, #3261, Corning, NY, USA) at a concentration of 10^4^ and 10^5^ cells per well or dish, respectively, in regular growth medium and cultured for up to 11 days. Conditioned Medium (CM) was isoDMEM-FCS harvested from primary cultures of fibroblasts derived from 13.5 days C57Bl/6 embryos and made senescent by gemcitabine treatment at 100 ng/ml for 7 days. The CM was filtered through 0.2 µm sized-pore filter and in some experiments concentrated with Amicon Ultra centrifugal filters with a cut-off of 50 kDa or 300 kDa.

### Identification of cancer cells in SSQ in low density 2D KC-DT66066 cell culture

Individual cancer cells pretreated with Torin-1 at medium cell density were followed from day 7 of the low FCS⟶ gem d3⟶ SSQi clonogenic assay by photographing them with a Olympus IX70 inverted phase contrast microscope using 20X LCPlan FI Ph1 and 10X UPlan F1 Ph1 objectives and a mounted Canon EOS 6D camera. Pictures (in jpg format; 5 to 12 pixels per microns) were manually analyzed using the polygonal enclosing tool of ImageJ. In spindle-shaped cells, the nuclear membrane was often indistinct, so we measured area of a nuclear region including nucleus and/or a perinuclear region enriched in vesicles, both exhibiting a specific granular texture and dark yellow-brown color (see Fig 4B). The mean value of the area of this nuclear region was close to that of the area of the nucleus measured in confluent KC-DT66066 cells in which nuclear membrane was unambiguously identified.

### Recombinant retrovirus

To downregulate *Beclin-1* expression we used recombinant lentiviral vectors, pLKO1-sh*Becn1*, that encode validated shRNA ^52, 53^ from the RNAi Consortium (clones TRCN0000087290 and TRCN0000087292 with target sequence 5’-GCGGGAGTATAGTGAGTTTAA-3’ and 5’-CGGACAGTTTGGCACAATCAA-3’, respectively, distributed by Sigma-Aldrich, Merck, France. pLKO1-sh*gfp* ^54^ was a gift from David Sabatini (Addgene plasmid # 30323; http://n2t.net/addgene:30323; RRID:Addgene_30323), pMRX-IP-GFP-LC3-RFP-LC3ΔG ^15^ a gift from Noboru Mizushima (Addgene plasmid # 84572; http://n2t.net/addgene:84572; RRID:Addgene_84572) and lenti-HA-mBec1 F121A ^22^ a gift from Congcong He (Addgene plasmid # 99506; http://n2t.net/addgene:99506; RRID:Addgene_99506). To produce retroviral particles, 2 µg of the plasmid carrying the relevant recombinant provirus was co-transfected with 2 µg of plasmid pMD2.G (a kind gift from D. Trono) and 2 µg of plasmid PVPackgagpol (#217566 from Stratagene, La Jolla, CA) or plasmid pCMV-ΔR8.74 (a gift from D. Trono) (for Moloney Murine Leukemia Virus and Human Immunodeficiency Virus derived retroviral backbone, respectively) in 90% confluent HEK293T cells plated in 6 cm diameter dishes using lipofectamine 3000 kit (12 µL of P3000 and 14 µL of Lipofectamine 3000 used). Supernatant were filtered through 0.4 µM filter and used to infect exponentially growing cells. Transduced cells were selected with puromycin (Calbiochem; 1 to 2 µg/ml) or directly used in some superinfection experiments with pMRX-IP-GFP-LC3-RFP-LC3ΔG.

### Flow cytometry analysis of autophagic flux

Cell transduced with MRX-IP-GFP-LC3-RFP-LC3ΔG vector were seeded at 4 x 10^5^ cells per 6 cm diameter dish (day 0), treated at day 1 as indicated and dissociated and harvested in cold culture medium at day 2 by strong pipetting after Tryp-EDTA-Pyr treatment, pelleted by centrifugation, resuspended in 500 µL of cold PBS 1X and fixed by adding 4.5 mL of 4 % paraformaldehyde in saline (sc-281692, Santa Cruz Biotechnology, France) and incubating at room temperature for 5 minutes. Then, cells were washed 2 times with PBS1X, suspended in 500 µL PBS 1X and kept at 4°C in dark until analysis. The GFP and RFP cell fluorescence was measured using a Fortessa Flow-cytometer (BD Biosciences) with excitation/emission at 488/515-545 and 561/600-620 nm respectively. For data analysis using FlowJo software (BD Biosciences), the RFP positive cell subpopulation was first selected to measure the GFP/RFP fluorescence ratio. The prior elimination of cell doublets did not have a significant impact on the results.

### mRNA isolation and RT-qPCR analysis

Cultured cells or spheroids were washed twice with cold PBS, and processed for total RNA isolation using the RNeasy Mini Kit (Qiagen, France) or Trizol reagent (Invitrogen, ThermoFisher Scientific, France). Tumor fragments were minced into fine pieces and lysed by vigorous shaking with steel balls in Trizol reagent. RT-qPCR experiments were performed as previously published ^8, 10^ using the LightCycler 480 SYBR Green I Master kit. *Hprt* was selected as the reference gene for normalization of data derived from independent cDNA synthesis reactions and normalization was further checked using others control gene such as *Sdha* or *Rplp0*. The expression level of each mRNA species was further normalized to that measured in the growing 2D-culture (KC-2D) set to 1. Oligonucleotide primers sequences along with a summary of data are shown in **S1 file**.

### Western blot analysis

Cells were seeded at 4 x 10^5^ cells per 6 cm diameter dish (day 0), treated at day 1 as indicated and lysed at day 2 or 3. For cell lysis, cells were washed two times with cold PBS1X and lysed in Laemli 2X lysis buffer supplemented with phosphatase inhibitors. About 50 µg protein lysate were subjected to SDS-PAGE and transferred to a polyvinylidene fluoride membrane (Hybond P; RPN303F from Amersham Pharmacia Biotech, England) that was blocked with 5% non-fat milk, in PBS1X containing 0.1% Tween-20 and probed with primary antibodies in the same buffer overnight at 4°C. Proteins were detected using horseradish peroxidase-conjugated secondary antibodies and detected using ECL (RPN2106) or ECL-select (RPN2235) reagents (Amersham, GE Healthcare, UK) and a cooled digital camera (LC33000, FujiFilm). Relative protein levels in each lane was measured using the Multi Gauge software (Fuji Microsoft, FujiFilm) associated to the digital camera device.

### Animal models

Six-week-old female *NOD.CB-17-Prkdc scid/Rj* (NOD-SCID) or C57Bl/6 J mice weighing 16–20 g were purchased from Janvier Laboratory (Le Genest Saint Isle, France). All mice were housed in ventilated cages with sterile food and water *ad libitum* throughout the study. The presence of subcutaneous tumors was carefully checked by palpation, light pinching and rolling of the skin between fingers to assess thickening of the skin and the presence of small hard lumps, which allowed to detect most of the tumors several days before they reach 1 mm^3^. Tumor size was measured two times a week with Vernier calipers. Tumor volumes were calculated using the following formula: (L * W^2^)/2, in which L represents the large diameter of the tumor, and W represents the small diameter. Mice were sacrificed when their tumor volume exceeded 1500 mm^3^ or when they showed signs of suffering.

### Experimental procedures for NOD-SCID mice

KC-DT66066 cells in SSQ were generated in two steps. 4 x 10^5^ cells were seeded in 6 cm diameter dish and treated the next day with 200 nM Torin1 for 1 day. Cells were then subcultured at low cell density in growth medium with 1.9% FCS plus insulin 10 μg/mL (to enhance survival of cells in SSQ) for 3 days and then subjected to gemcitabine selection for 5 days in regular medium supplemented with gemcitabine at 30 ng/mL. Cells in SSQ were then harvested in DMEM with 10% fetal calf serum after 5 min treatment with Tryp-EDTA-Pyr solution warmed to 37°C, washed two times with cold DMEM without serum or antibiotics, resuspended in a small volume of DMEM and counted. Exponentially growing cells were harvested from 6 cm diameter dishes seeded with 4 x 10^5^ cells two days before and harvested as cells in SSQ in DMEM. Harvested cells were injected subcutaneously in the backs of mice. Gemcitabine at 50 mg/kg was injected intraperitoneally on day 4 and 11 for the Ssq/gem group, day 0 referring to the day of cell injection.

### Experimental procedures for C57Bl/6 mice

Cells were seeded at 4 x 10^5^ per 10 cm diameter plate and then cultured for 4 or 5 days with a growth medium change on the third day. Cells were harvested by a 3 min treatment with Tryp-EDTA-Pyr, washed twice with growth medium without serum or antibiotics, filtered through a 70 μm cell strainer and 200 μl of a cell suspension at 5 x 10^6^ cells/mL subcutaneously injected (day 0) into the back of C57Bl/6 mice. Cyclosporin A (Sandoz, France) was administered orally at a dose of 80 mg/kg at days −2, 0, 4, 7 and 11.

### Analysis of patients’ records

The anonymized records from 144 patients with pancreatic adenocarcinoma (PAAD) included in the TGCA-PAAD project and with a follow up and a transcriptome profiling of their primary tumors by Illumina sequencing were downloaded from the Genomic Data Commons website as clinical files and FPKM transcriptome-sequencing files. For gene expression analysis, genes counts from the FPKM files were assembled into a count matrix K_ij_ (for gene i in tumor sample j) after filtering genes with zero geometric mean. Gene counts were normalized by calculating scaling factors analogous to those calculated in the DESeq package ^55, 56^: 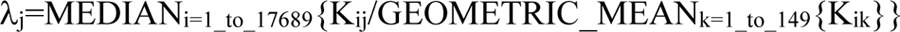. The normalized FPKM value K_ij_/α_j_ was used as a measure of the expression level of gene i in tumor j. In the case of 5 patients for whom there were 2 sequencing files, we took the geometric mean value of normalized gene counts. This normalization introduced in most cases only a small correction (the average value of α_j_ was α=1.01 ± 0.12, mean ± s.d.; n=149) and was validated *a posteriori* by the strong self-consistency of the results obtained. The cutoff value θ used to categorize patients in cohorts with a low or high level of expression of a given gene in their tumor, was chosen to maximize the statistical significance of the difference in survival between the two cohorts of patients. For *BECN1* mRNA levels (8.56 ± 1.83 FPKM, mean ± s.d., range: 4.2-18.0, n=124), a cutoff value of 8.895 FPKM was used throughout this study. For *CCNA2* mRNA levels (4.76 ± 2.86 FPKM, mean ± s.d., range: 1.12-16.44, n=124) a cutoff value of 5.66 FPKM was used throughout this study (see also **S2 file**).

The clinical data were verified by cross-referencing the various clinical files for the treatment status and for the duration of patient survival without local or distant tumor recurrence. Only the records of 124 of the 144 patients contained relevant indications on local recurrence and the status of distant metastasis. They include 109 patients with a pancreas adenocarcinoma of the ductal subtype, 6 with a Not Otherwise Stated subtype, 3 with a poorly differentiated and 1 with a well differentiated subtype, 3 with mixed phenotypes (ductal and micropapillary or neuroendocrine subtypes) and 2 with an invasive phenotype. The “Initial treatment” corresponded to removal of the primary tumor by partial or total pancreatectomy. Adjuvant chemotherapy was recorded for 78% of patients (the treatment status was not specified for the remaining patients) and this percentage increased to 89% for patients with distant metastasis-free survival >360 day. When a patient had a local recurrence or a new cancer type, he was censored at the time of that diagnostic in the Kaplan-Meier analysis for distant metastasis-free survival. Data were processed and analyzed using Excel (version 16; Microsoft, USA) upgraded with Real Statistics add-ins (Zaiontz, C., 2020, Real Statistics Using Excel.at www.real-statistics.com) and GraphPad Prism version 5.00 (San Diego, California, USA). Enrichment analysis was performed with Gene Set Enrichment Analysis software (GSEA v4.2.0 for Windows and Mac) with default settings and the msigdb.v7.4.symbols.gmt dataset available from the Broad Institute. Hierarchical gene clustering was performed using Morpheus software (https://software.broadinstitute.org/morpheus). The references of downloaded files and a synthetic summary of clinical data are given in **S2 file**. Normalized sequencing data are provided in **S3 file**.

### Statistics

In all graphs, each point corresponds to an independent measurement, the labels above the data points indicate the average values and error bars are standard deviation unless otherwise indicated. Significance was measured using a two-tailed Student’s t-test unless otherwise specified. Log-ranked (Mantel-Cox) and Fisher’s exact probability test were performed using GraphPad Prism version 5.00 or 8.00 (San Diego, California, USA). Correlations were calculated using Excel software (Microsoft Standard Office 2016) and their statistical significance calculated on the Vassar web site (http://vassarstats.net). Multivariate Cox regression analysis was performed using Excel (version 16; Microsoft, USA) upgraded with Real Statistics add-ins (Zaiontz, C., 2020, Real Statistics Using Excel.at www.real-statistics.com). The number of * indicates the statistical significance of the difference between the indicated mean values with * for p<0.05, ** for p<10^-2^, *** for p<10^-3^, **** for p<10^-4^, ***** for p<10^-5^ and ******* for p<10^-6^ and the same for #.

## Supporting information

Appendices section Fig S1-S4

Appendices section File S1

Appendices section File S2

Appendices section Fig S3

## Declarations and ethics satements

### Ethical approval

The animal protocol used in this study was reviewed and approved by the local ethics committee (Comité d’Ethique en Expérimentation Animale of University of Paris; approved protocol CEEA 18-088 submitted by C. Nicco and T. Tchenio). Animals received humane care in compliance with the guidelines implemented at our institution (INSERM and University Paris Descartes).

### Data availability statement for Share Upon Reasonable Request Data

The authors declare that all data supporting the findings of this study are available within the article and its appendices, or available from the corresponding author upon reasonable request.

### Author contributions

Conceptualization: TT. Supervision: CN, FB, TT. Methodology: TT. Investigation: JG provided murine pancreas cancer cell lines derived from corresponding transgenic mice; MH carried out counting of colonies from clonogenic assays and plasmids constructions in studies with prostate cancer cells. TT performed all cell culture, flow cytometry, qPCR, Western Blot experiments and analysis of patients’ records; CN and MT carried out animal experiments. Formal analysis: T.T; Analysis and Visualization of Data: FB, CN, TT. Writing original draft: TT. Review and Editing: CN, FB, FD, FLT, JG, LD and TT.

## Appendices

**Fig S1: Selection of cancer cells in SSQ with anticancer agents in pancreas and prostate cancer cell lines cultured at clonogenic cell density. A)** Delayed addition of SSQi after gemcitabine selection does not impact recovery of growing cell clones. Pictures of KC-DT66066 cell culture dishes from a representative experiment (of 3) fixed and stained after cell culture as indicated. **B)** Selection of cancer cells in SSQ with gemcitabine using indicated pancreas cancer cell lines derived from KC and KPC mouse models. **C)** Representative pictures showing the formation of spheroid-like structures in KC-A338 but not KC-DT66066 cells cultured in 2D. Scaling bar indicates 1 mm. **D)** Comparison of the selection of KC-DT66066 cell in SSQ with 100 nM nocodazole (noco), 30 ng/mL gemcitabine (gem) and 100 ng/mL vincristine (vin). **E)** Selection of SSQ in LNCaP* cells using doxorubicine (dox) and Docetaxel (Tx) at the indicated concentrations. Note that cells were selected from day 0 to day 8 (d0-d8), washed and further cultured for around 20 days in regular growth medium supplemented with none or SSQi. Labels indicates the percentage of seeded cells having formed a cell clone after supplementation with SSQi. **F)** clonogenic assays for testing induction of SSQ in KC-DT66066 cell populations generated after one SSQ induction/selection/reversal cycle using gemcitabine (n=2) or nocodazole (n=2) to select cells in SSQ. Cell populations were derived from experiments depicted in Fig S1D. Stars indicate statistical significance of cloning efficiency differences between cells untreated and treated with SSQi after drug selection.

**Fig S2: SSQ induction is regulated by mTOR and Beclin-1 but not autophagic flux. A)** Increased frequency of C4-2 cells in SSQ induced by treatment with 200 nM Torin1 from day 1 to day 3 (torin1 d1-d3; cells were seeded at low cell density on day 0). **B**) Increased frequency of KC-DT66066 cells in SSQ induced by treatment of cells with 5 nM rapamycin from day 0 to day 3 (rap d0-d3; cells were seeded at low cell density on day 0). **C)** Downregulation of Beclin-1 expression significantly decreases frequency of SSQ cells in KC-A338 cell populations. Cell populations transduced with sh*gfp* (control) or sh*Becn1* expressing vectors were assayed with the gem d0 clonogenic assay as described in Fig. 1. **D**) No significant effect of autophagy inhibitor bafilomycin A1 on SSQ induction by Torin1. Native KC-DT66066 cells were pretreated at medium cell density with none (no pretreatment), 100 nM bafilomycin A1 (baf1 pre-treatment), 200 nM Torin1 (tor1 pretreatment), or bafilomycin A1 + Torin1 (baf1+tor1 pretreatment) for 1 day prior to cell seeding at low cell density for assaying SSQ induction. Note that bafilomycin A1 induced delayed toxicity (compare CE in the “none” clonogenic assays for the different pretreatment groups) that at least partially explained the non-significant difference in SSQ induction between the tor1 and baf1+tor1 pretreatment groups. **E)** Same as D) but performed in human C4-2 cells. Cells were seeded at low density on day 0 and treated as indicated from day 2 to 3 before gemcitabine selection. It is one of 2 experiments done with either Torin1 or rapamycin. **F)** Western Blot analysis of KC-DT66066 transduced with empty (control) or HA-*Becn1^F121A^*-expressing retroviral vectors. Note the higher molecular weight of HA-BECN1*^F121A^*compared to wild type (wt) BECN1. **G)** Comparison of autophagic flux between control and HA-*Becn1^F121A^*-transduced cell populations cultured at medium cell density and treated as indicated for one day before cytometric analysis to measure gfp/mRFP fluorescence ratio. 4 independent cell populations were generated by superinfection with MRX-IP-GFP-LC3-RFP-LC3ΔG retroviral expression vector and assayed for autophagic flux 4 days later. **H)** Similar rate of SSQ induction in control or HA-*Becn1^F121A^*-transduced cell populations as assayed with the indicated clonogenic assays. Labels indicates the percentage of seeded cells that formed a cell clone after SSQi supplementation.

**Fig S3: Tumorigenicity of KC cells injected subcutaneously in mice. A)** Tumor growth curves in allogenic C57Bl/6 mice injected subcutaneously with 10^6^ KCsh*gfp* or KCsh*Becn1* cells. The errors bars correspond to the standard error of the mean. * indicates statistically significant differences (p<0.05) between the 2 groups of mice. **B)** Same as A but mice were treated once with gemcitabine (100 mg/kg i.p.) on day 4 after cell injection. **C**) Comparison of cell proliferation between KC*sghgfp* and KC*shbecn1* cell populations (3 independent pairs) seeded à 4×10^5^ cells in 6 cm diameter dish and cultured for 2 days in regular growth medium before cell counting. **D**) Tumor growth curves of individual NOD-SCID mice injected with 10^2^ and 10^3^ dissociated KC-DT66066 cells harvested from growing 2D cell-cultures. **E**) Comparison of mRNA species levels in growing tumors derived from indolent tumors (n=5 in blue) to those measured in Exp* group tumors (n=4 in red) and in KC-DT66066 cells cultured in 2D at medium cell density (in black, normalized to 1 as in Figure 2C and File S1). Labels in black indicate number of independent experiments for growing KC-DT66066 cells in 2D-culture. * indicates the statistical significance of the differences between tumors and KC-DT66066 cells in 2D-culture.

**Fig S4: Beclin-1 and Cyclin A2 transcript levels in the primary tumor of PAAD patients are predictive of early and late metastasis. A)** Kaplan-Meier curves for distant metastasis-free survival restricted to PAAD patients with T3-N1-M0/Mx clinical grade and according to the level of Beclin-1 transcripts in their primary tumor. Statistical difference (p) in survival and hazard ratio (HR) are indicated. B) Same as panel A but also restricted to patients with distant metastasis-free survival time greater than 360 days. C) Miscellaneous gene set enrichments also found in primary tumors of PAAD patients with distant metastasis-free survival time less than 360 days after initial treatment (n=60) *versus* greater than 360 days (n=64). Normalized enrichment score (NES), its statistical significance (p) and its false discovery rate (FDR) are indicated. D) Kaplan-Meier curves for distant metastasis-free survival restricted to PAAD patients with T3-N1-M0/Mx clinical grade and according to the level of Cyclin A2 (CCNA2) transcripts in their primary tumor. E) Gene set enrichment found in the primary tumors of PAAD patients with high level (n=31) *versus* low level (n=93) of CCNA2 transcripts. F) Pearson’s correlation coefficient between Beclin-1 or Cyclin A2 transcript levels and those of cancer, mesenchymal, endothelial, and immune cell marker genes in primary tumors of PAAD patients (n=124). The stars indicate the statistical significance of the correlation.

**S1 file:** Sequences of primers used for RT-qPCR analysis and synthetic summary of RT-qPCR expression data.

**S2 file:** Synthetic summary of clinical and sequencing data including references of clinical and sequencing files retrieved from the Genomic Data Commons website.

**S3 file:** Genes count matrix derived from the 149 FPKM sequencing files (retrieved from the Genomic Data Commons website) and normalized by scaling factors 1χ_ij_. See the Materials and Methods section for further explanations.

