## Supplementary figures and images for "Mechanistic target of rapamycin (mTOR) regulates self-sustained quiescence, tumor indolence and late clinical metastasis in a Beclin-1-dependent manner"

### Appendices section Fig S1-S4

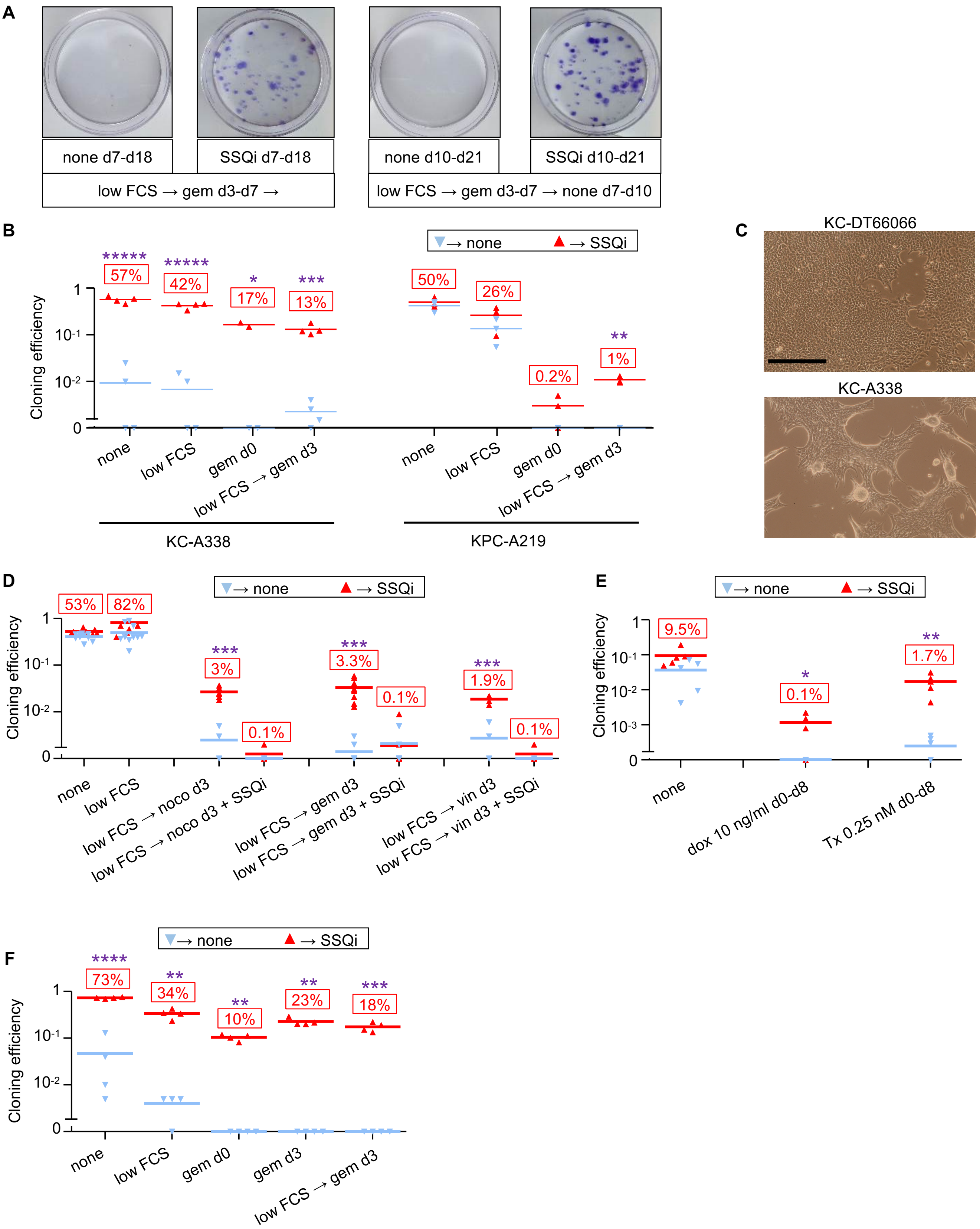

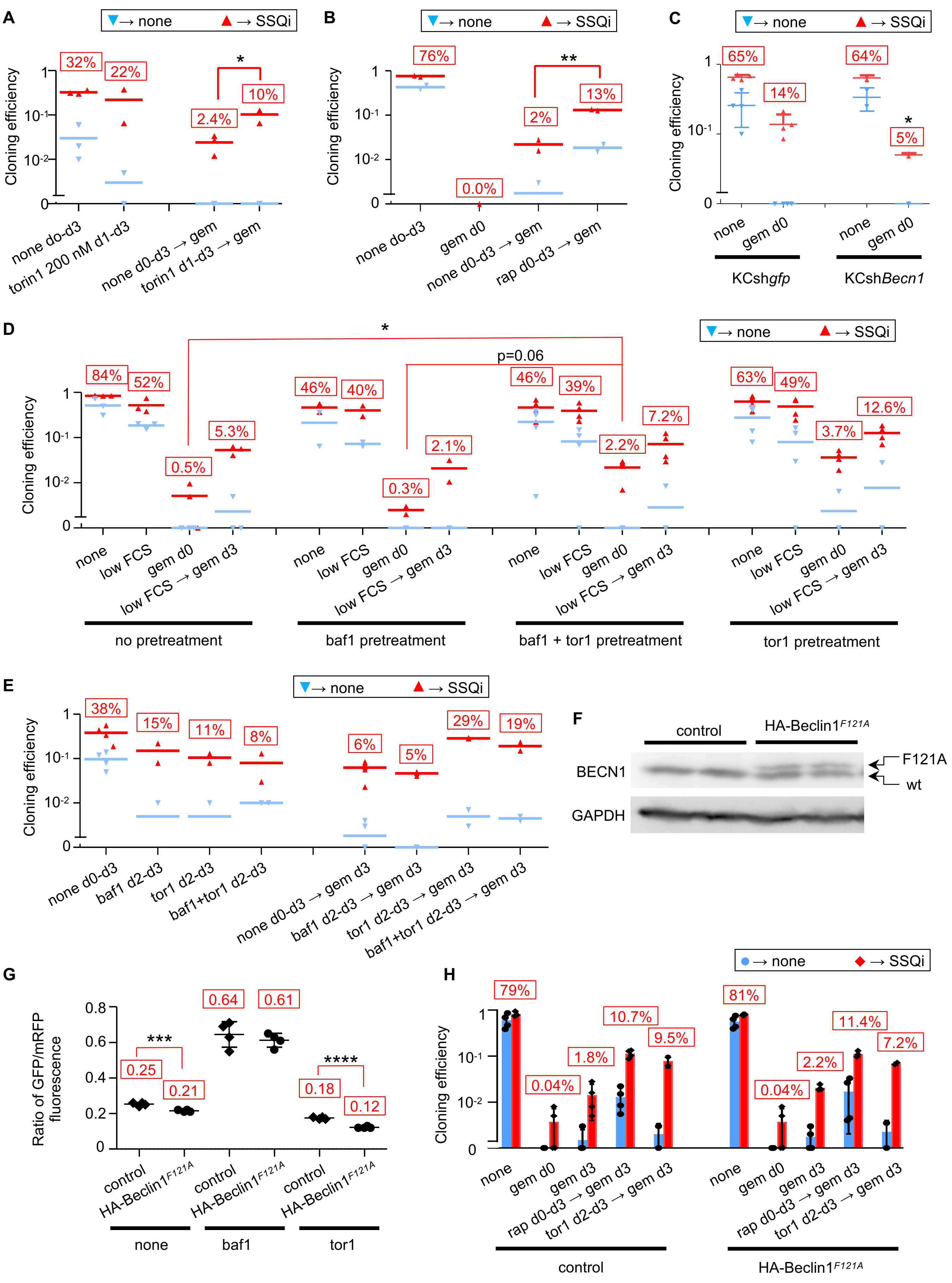

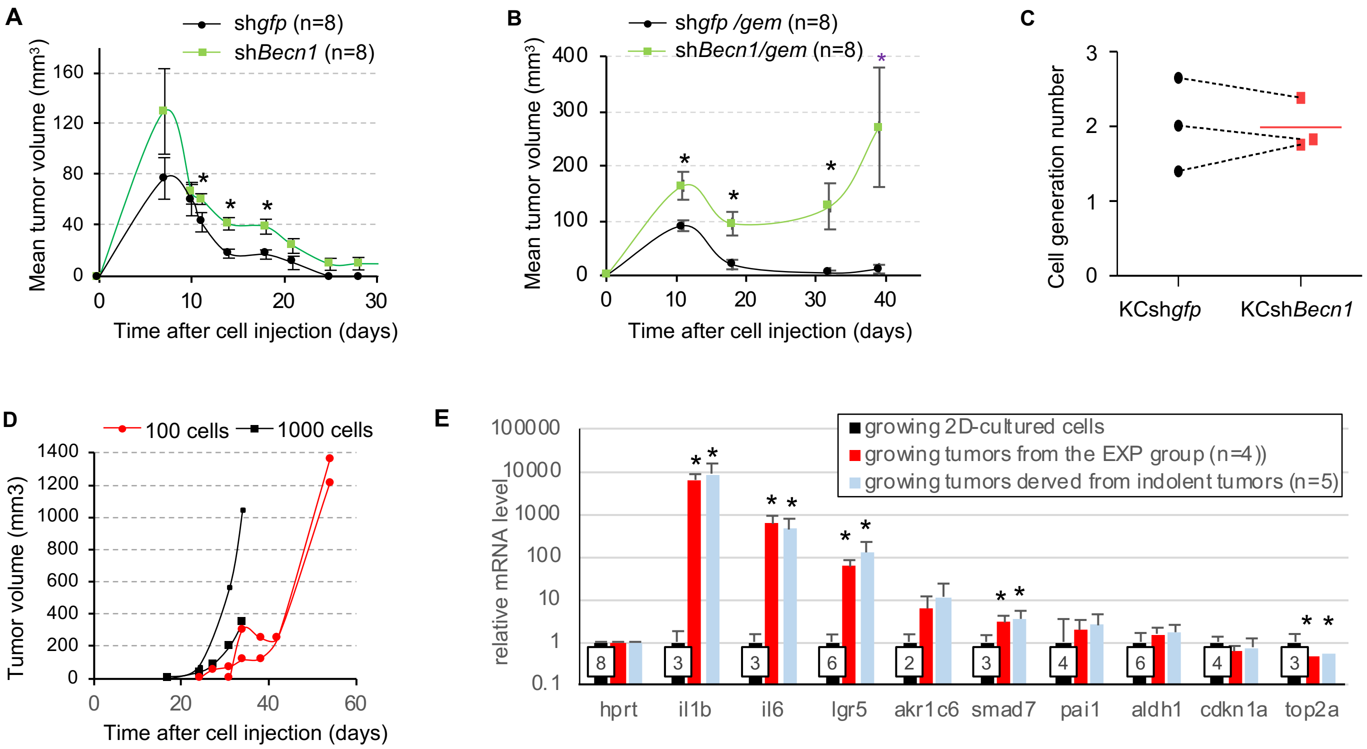

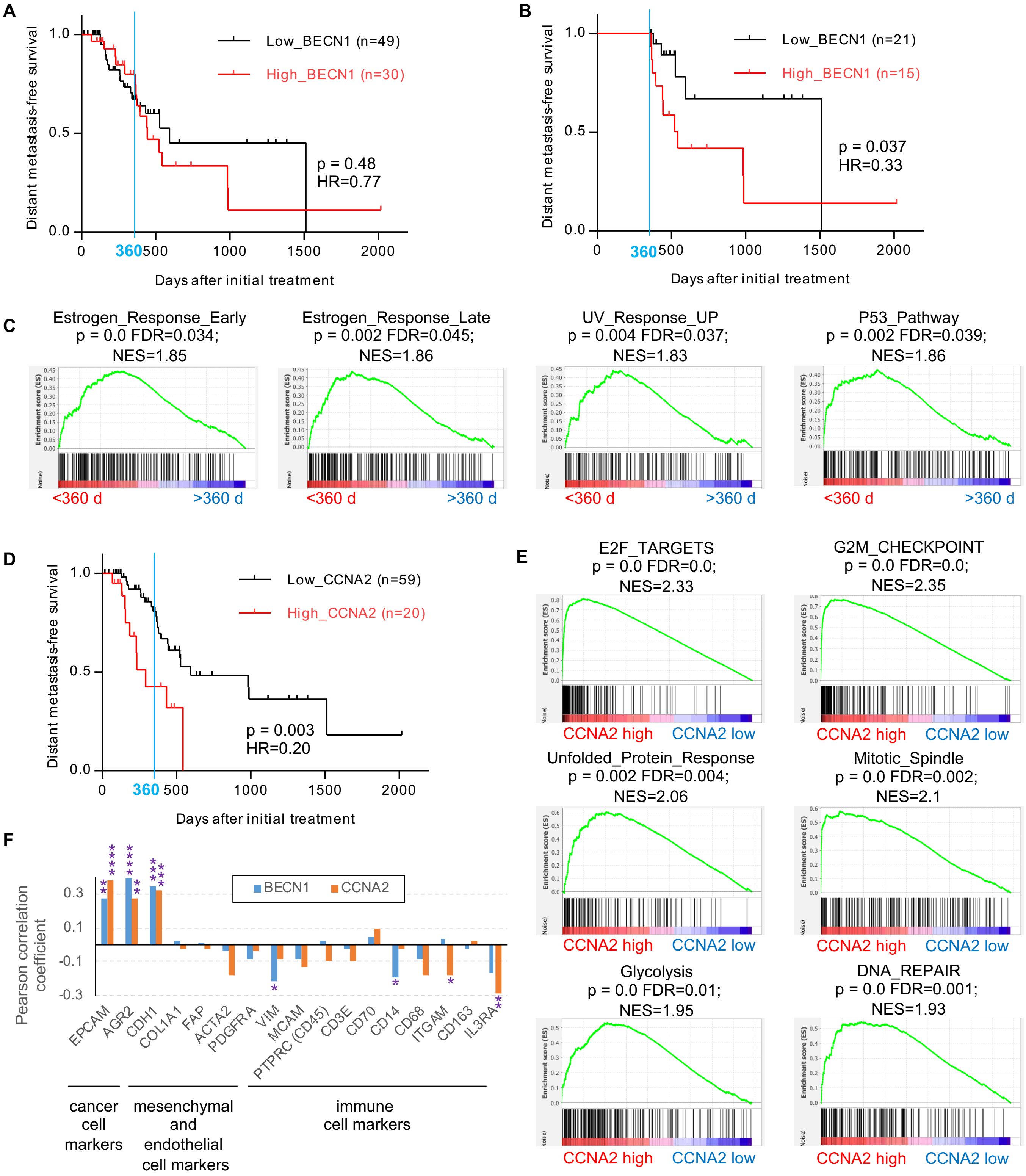
